# Vectorized instructive signals in cortical dendrites during a brain-computer interface task

**DOI:** 10.1101/2023.11.03.565534

**Authors:** Valerio Francioni, Vincent D. Tang, Enrique H.S. Toloza, Norma J. Brown, Mark T. Harnett

## Abstract

Vectorization of teaching signals is a key element of virtually all modern machine learning algorithms, including backpropagation, target propagation and reinforcement learning. Vectorization allows a scalable and computationally efficient solution to the credit assignment problem by tailoring instructive signals to individual neurons. Recent theoretical models have suggested that neural circuits could implement single-phase vectorized learning at the cellular level by processing feedforward and feedback information streams in separate dendritic compartments^1–5^. This presents a compelling, but untested, hypothesis for how cortical circuits could solve credit assignment in the brain. We leveraged a neurofeedback brain-computer interface (BCI) task with an experimenter-defined reward function to test for vectorized instructive signals in dendrites. We trained mice to modulate the activity of two spatially intermingled populations (4 or 5 neurons each) of layer 5 pyramidal neurons in the retrosplenial cortex to rotate a visual grating towards a target orientation while we recorded GCaMP activity from somas and corresponding distal apical dendrites. We observed that the relative magnitudes of somatic versus dendritic signals could be predicted using the activity of the surrounding network and contained information about task-related variables that could serve as instructive signals, including reward and error. The signs of these putative teaching signals both depended on the causal role of individual neurons in the task and predicted changes in overall activity over the course of learning. Furthermore, targeted optogenetic perturbation of these signals disrupted learning. These results provide the first biological evidence of a vectorized instructive signal in the brain, implemented via semi-independent computation in cortical dendrites, unveiling a potential mechanism for solving credit assignment in the brain.

## Dendrites as a biological substrate for credit assignment in the brain

Learning is the product of changes in the strength of synaptic connections between neurons^6–13^. Synaptic modifications can have difficult-to-predict effects on network output, particularly in complex hierarchical networks like the brain. The challenge of determining how individual synapses should be altered to improve task performance is known as the credit assignment problem^14–18^. While this problem is effectively solved in artificial neural networks (ANNs) by the backpropagation-of-error algorithm^19^, how credit assignment is solved in the brain remains unknown^14,15^.

Recent theoretical work has proposed several models by which biological circuits could solve credit assignment, including target learning and backpropagation-like algorithms^1–5,20,21^. Central to both artificial and biologically-inspired solutions to credit assignment is the vectorization of instructive signals, as opposed to the broadcasting of a single scalar teaching signal^14^. Effective learning requires, in addition to vectorization, instructive signals to be separable from feedforward inputs to prevent interference^15^. In ANNs, this is achieved via temporal separation, which has long been thought to be biologically-implausible. One hypothesis is that in cortex, credit-related information is spatially, rather than temporally, segregated in the apical dendrites of pyramidal neurons^15^. This aligns with anatomical and circuit evidence that feedforward inputs are received perisomatically and feedback inputs are received in the distal dendrites^22–30^. However, direct evidence regarding the subcellular mechanisms of credit assignment is lacking.

Vectorized teaching signals at the dendritic level should meet four experimentally testable criteria. First, dendritic activity should contain information not present in somatic activity alone (while somas could theoretically transmit gradients using qualitatively different spiking patterns^2,4,31^, the cable properties of dendrites predict some level of independence between somatic and dendritic activity). Second, dendritic activity should encode information about task performance that could serve as instructive signals, such as reward and error-representations. Third, dendritic activity should reflect the contribution of that neuron to task performance (i.e., the reward function). Fourth, disrupting vectorized instructive dendritic signals should impair learning.

## Specifying a reward function using a brain-computer interface task

Evaluating credit assignment in biological neural networks has thus far proven impossible^14,15^. Teaching signals can only be defined relative to a reward function that maps neural activity to task performance. It is unclear if such functions are explicitly represented in the brain. Even if they are, experimenters are blind to their specific formulation in terms of neural activity^15^. Neurofeedback brain-computer interface (BCI) tasks present a potential solution to this problem by directly coupling neural activity to task performance, thereby allowing the experimenter to specify the reward function to be optimized^14,20,21^. Previous studies have shown that mice are able to learn BCI tasks using a variety of feedback stimuli and brain areas and that learning induces changes in the activity of the neurons controlling the BCI, including in the hippocampus and various sensory and motor cortices^32–38^. Here, we leveraged a visually guided neurofeedback BCI task in cortical pyramidal neurons to test subcellular mechanisms for error and reward-related signaling (Fig. 1a, b and c, and Extended Data Fig. 1 and Extended Data Fig. 2). We trained head-fixed mice under a 2-photon microscope to control the activity of two spatially intermingled sets of GCaMP7f-labeled layer 5 (L5) pyramidal neurons (PNs), in the retrosplenial cortex (RSC), designated P+ and P-(see methods Extended Data Fig. 3 and Extended Data Fig. 6b for selection criteria). The difference in mean somatic GCaMP activity of P+ versus P-neurons was coupled to rotation of a visual grating relative to a rewarded target angle^32–35,37,38^ (Fig. 1d, e, and f and Extended Data Fig. 1). We selected RSC due to the optical accessibility of layer 5 and previous demonstration of independent dendritic events in this area^39^. We recorded GCaMP activity at 15 Hz in the proximal trunk dendrite as a proxy for somatic activity; this allowed imaging of many neurons while reducing signal contamination due to the more precise spatial footprint and faster signal kinetics of the apical trunk^40–42^. We measured task performance with two metrics: accuracy, which represented the fraction of rewarded trials; and speed, which represented the numbers of rewards obtained per minute. Mice (n = 6) learned the task by both metrics (Fig. 1g, Extended Data Fig. 4 and Extended Data Fig. 5).

**Figure 1:**
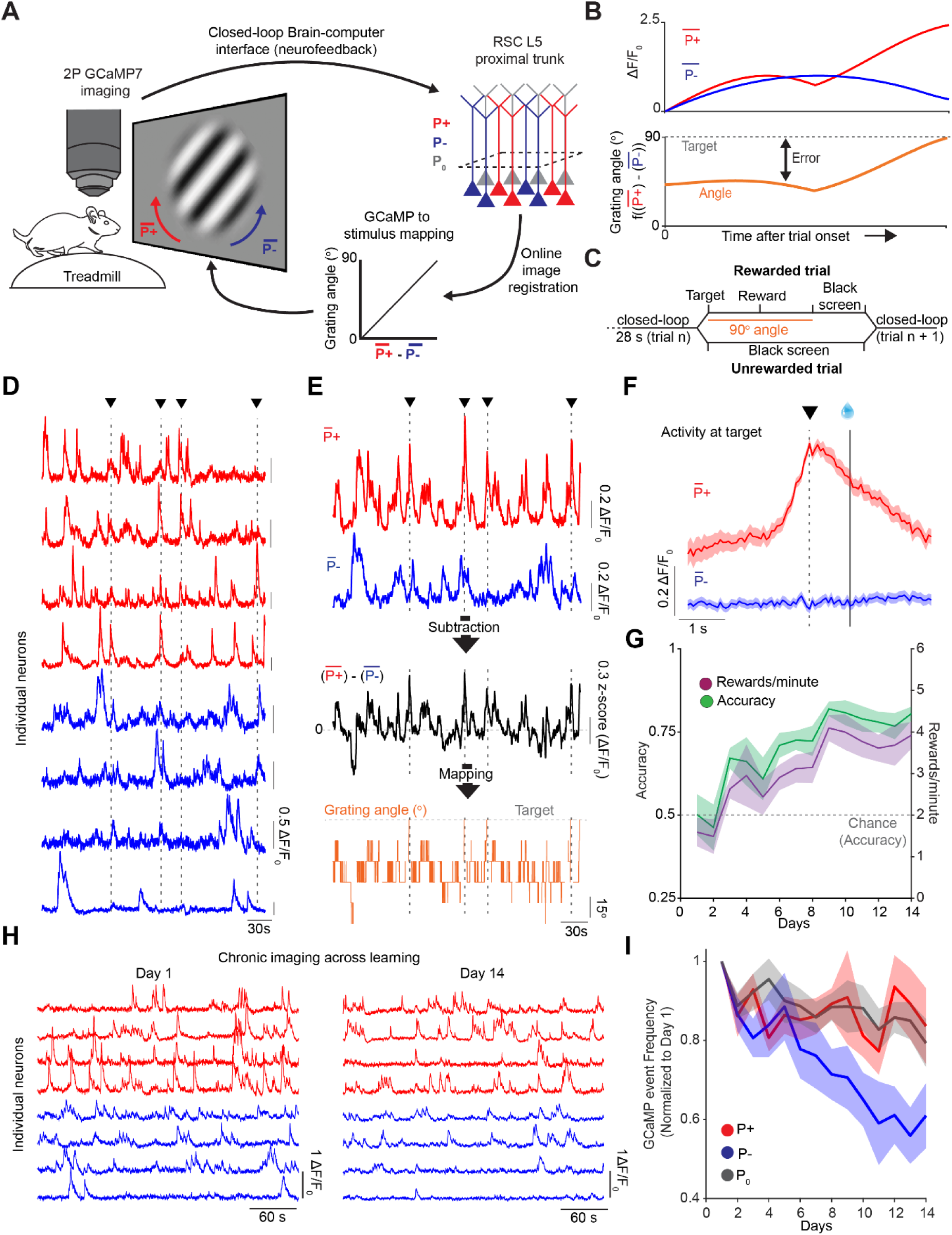
Mice learn a neurofeedback BCI task through the differential regulation of P+ and P-neurons. **a,** Schematic of the BCI setup. Mice were head-fixed and imaged under a 2-photon microscope and free to run on a cylindrical treadmill. Two user-defined populations of GCaMP7f-labeled layer 5 pyramidal neurons in retrosplenial cortex (RSC) were imaged at the proximal apical trunk: P+ (in red) and P-(in blue) were selected to control the rotation of a Gabor patch. P_0_ neurons were designated as all other neurons in the field of view. Single frames were online-registered (motion-corrected). Activity in P+ neurons rotated the patch clockwise, towards the target angle of 90-degree. Activity in P-neurons rotated the Gabor patch stimulus counter-clockwise, towards a 0-degree angle. **b,** Schematic of the mapping between P+ and P-activity, stimulus angle, target activity and error. Error was the distance between current and target activation. The angle represents a binned (7 bins, 15 degrees apart, from 0 to 90 degrees) linear mapping between the mean activity in P+ neurons minus the activity in P-neurons. **c,** Trial structure: mice had 28 seconds to reach target activity and receive a reward delivered 1 second later. In successful trials, the 90-degree Gabor patch was shown for 2 seconds, followed by 1 second of black screen presentation. In unsuccessful trials, a 3 second black screen was presented before the onset of the next trial. **d,** ΔF/F_0_ traces as recorded live for P+ (in red) and P-(in blue) neurons. Vertical dashed lines and triangles represent timepoints where the animal reached target activity. **e,** Mean activity for the red (P+) and blue (P-) traces shown in d. Black trace shows the arithmetic subtraction of P+ and P- neurons (z-scored). Orange trace shows the corresponding visual stimulus angle as presented to the mouse. **f,** Mean ΔF/F_0_ for P+ and P- activity aligned to the time in which the animal reached target activity (dashed, vertical line and black triangle) for the session highlighted in d and e. Reward was delivered 1 second later (solid vertical line with water reward). Shaded areas are standard error of the mean (SEM). **g,** Mean performance over days quantified as the fraction of successful trials over the total number of trials in green, and as the number of rewards per minute in purple (One-way repeated measures ANOVA, p = 5e^-4^ and p = 0.002 for accuracy and rewards/minute, respectively. n = 6 mice). Dashed horizontal red line represents chance level for accuracy performance (see Methods). **h,** ΔF/F_0_ traces for the same P+ and P- neurons on training day 1 and training day 14. **i,** Calcium transient frequency for P+, P- and P_0_ neurons (in red, blue, and black, respectively) across the 14 days of training normalized to the activity on day 1. All neurons were tracked over the full 14 days of imaging. (Two-way repeated measures ANOVA, p = 0.012, p = 0.004 and p = 9.3e^-4^ for the effect of population identity, days and an interaction between these 2. After Tukey’s multiple comparison, p = 0.027, p = 0.95 and p = 0.01 for P+ vs. P-, P+ vs P_0_ and P- vs P_0_ neurons, respectively. n = 6 mice).

We compared activity levels of P+ and P- populations, as well as the population of surrounding neurons not directly involved in the rotation of the stimulus (termed P_0_), across days of task performance. We imaged the same neurons longitudinally throughout all experiments. We found that learning was accompanied by the differential regulation in the activity of P+ and P- neurons over days (Fig. 1h, i), with P+ neurons maintaining their activity levels while P- neurons were downregulated While, on average, changes in activity in P_0_ neurons resembled changes in P+ neurons (Fig. 1i), selecting the subpopulation of P_0_ neuron with matching activity levels of P+ and P- neurons on day 1 revealed that changes in activity in P_0_ neurons fell in between P+ and P- neurons (Extended Data Fig. 6). As the most active neurons on day 1 were also those most strongly downregulated (Extended Data Fig. 6b), our results are consistent with a model of learning by sparsification, an energy-efficient solution to the task^43^. Increases in task performance were not correlated with changes in locomotion across days (Extended Data Fig. 7). Moreover, the P+ and P- populations were spatially intermingled, and had the same GCaMP transient frequency on day 1 (Extended Data Fig. 3 and Extended Data Fig. 6a), ruling out the possibility of learning the task by simply engaging a non-specific gain modulation mechanism.

## Dendrites contain information not found in their somas

To determine whether apical dendritic activity contained information not encoded in parent somatic activity alone, we used an electrically tunable lens (ETL) to semi-simultaneously (15 Hz per plane) record activity in proximal and distal trunk dendrites across learning (Fig. 2a). We paired proximal and distal dendrites based on the Pearson correlation of their GCaMP signals, thresholded at r = 0.6 as in previous studies^40–42^. Previous work in brain slices demonstrated that dendritic GCaMP signals are larger when current is injected in the distal trunk and smaller when current is injected at the soma^40^ (controlling for the same number of triggered corresponding action potentials). This indicates that differences in somatic versus dendritic magnitude for coincident GCaMP events reflect the spatial bias of the different inputs that target these two compartments. To estimate the magnitude of somatic and dendritic events, we first deconvolved the GCaMP traces of somas and dendrites using CASCADE^44^. Deconvolution allowed us to correct for the well-described problem of different signal kinetics across dendritic compartments^45^. Next, we utilized an area-under-the-curve approach to quantify the magnitude of individual transients (all main results were also validated using a ΔF/F_0_-based approach to estimation of transient’s magnitude, see Methods and Extended Data Fig. 8) and defined events as coincident whenever they occurred within 500 ms of each other. Since these coincident events represent the vast majority of GCaMP transients^39–42,45–51^, we focused all subsequent analysis on events for which a transient was detected in both compartments.

**Figure 2:**
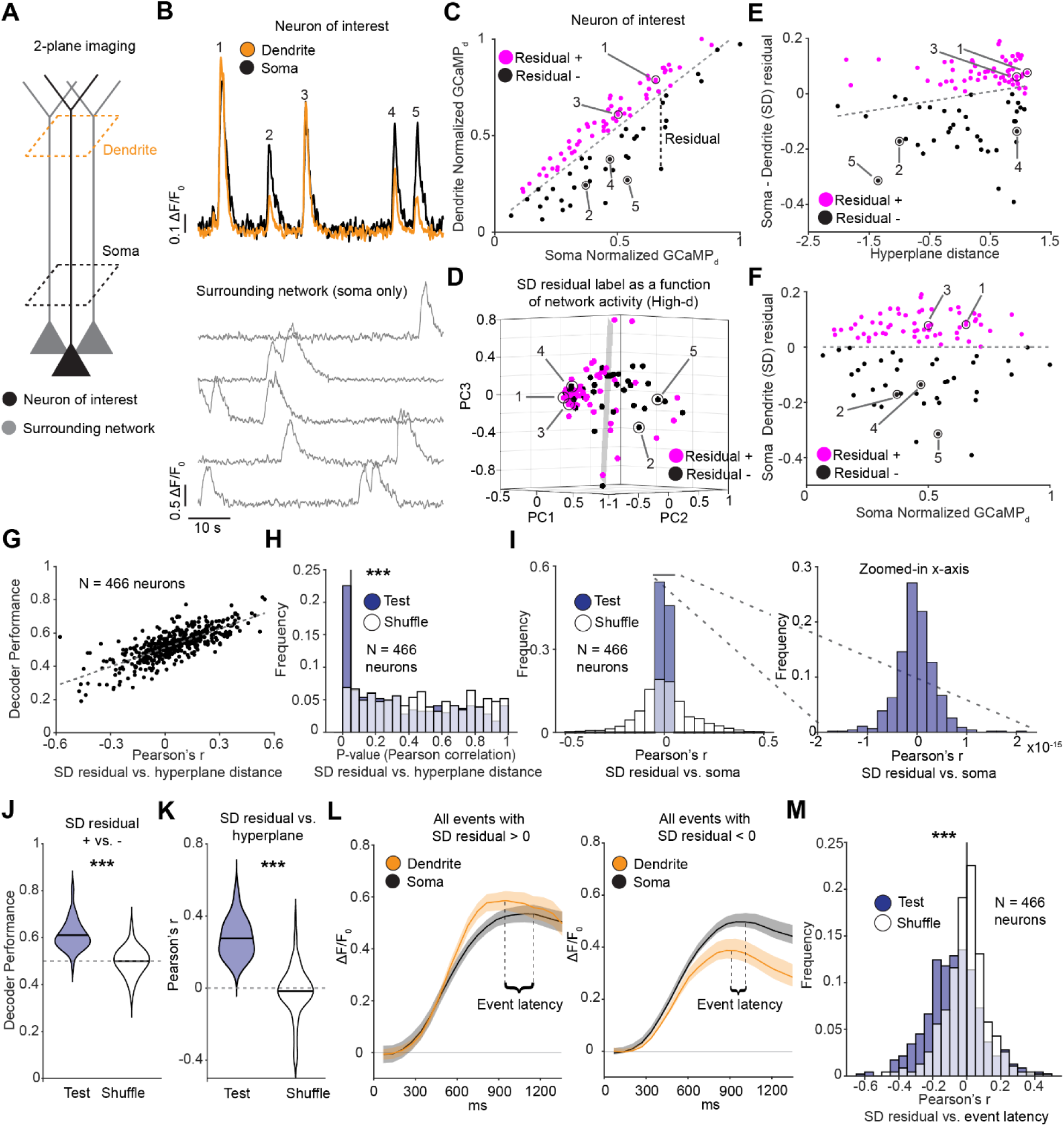
Differences in somatic and dendritic magnitudes for coincident events are predicted by local network dynamics. **a,** Schematic of two-plane 2-photon calcium imaging of a network of neurons at the proximal and distal trunk. **b,** ΔF/F_0_ traces recorded simultaneously in the soma (black) and dendrite (orange) for a single neuron of interest (top, P+ and P-neurons across days 1-14) and corresponding activity in 5 surrounding neurons. Numbers 1-5 indicate identified GCaMP events. **c,** Relationship between somatic and dendritic transients’ integral for the example neuron shown in b. Datapoints represent individual events simultaneously detected in soma and dendrite (see Methods). A least-squares linear model (dashed grey line) defined events as dendritically amplified (in magenta, residual +) versus dendritically attenuated (in black, residual -). Events (1-5) correspond to the transients shown in b. **d,** For each coincident event in the neuron of interest shown in b and c, we estimated the network activity vector in the 2 seconds before, using all other neurons in the field of view. Here, network activity vector was projected onto the first 3 principal components for visualization only. The shaded black hyperplane represents the decision boundary for binary classification (dendritically amplified versus dendritically attenuated) calculated using a linear SVM. Events 1-5 correspond to the network activity vector associated with transients 1-5 shown in b and c. **e,** The relationship between SD residuals estimated in c and the distance from the decision boundary (hyperplane distance) estimated in d for all coincident somato-dendritic events in the neuron of interest. Events 1-5 correspond to those shown in b, c and d. The dashed grey line represents the least-squares best-fit line. To maintain visual consistency with d, the distance from the hyperplane was estimated on the first 3 principal components. This is for visualization only. **f,** The relationship between SD residual as estimated in c, and somatic event magnitude. Highlighted events 1-5 correspond to those shown in b, c, d and e. Dashed grey line represents the least-squares best fit line. **g,** Decoder performance as a function of the correlation between SD residuals and hyperplane distance (Pearson’s r = 0.74; p-value = 1.4e^-84^, n = 466 neurons). Datapoints represents individual neurons. **h,** Distribution of p-values for test data and a control randomly shuffled distribution, testing the correlation between SD residuals and distance from the hyperplane (or classification confidence, Wilcoxon signed rank test = p = 1.3e^-9^. N = 466 neurons) as estimated in e. **i,** Left panel, for all neurons, the Pearson’s r for somato-dendritic residuals and somatic event magnitude as characterized in f. The residual-based approach perfectly decorrelates SD residual from somatic activity alone. Right panel, for test data, a zoomed in version of the same histogram shown in the left panel. Note the 10^-15^ scale on the x-axis. **j,** Decoding performance for neurons with a statistically significant correlation between SD residual and distance from the hyperplane (paired t-test, p = 8.6e^-9^. Mean = 0.61 and 0.50; SEM = 0.006 and 0.007; n = 82). Dashed grey line indicates chance level. **k,** Pearson’s r for neurons with a statistically-significant correlation between SD residual and the distance between population vector and hyperplane (paired t-test, p = 3.35e^-25^. Mean = 0.28 and-7.2e^-4^; SEM = 0.01 and 0.01; n = 82). **l,** Mean ΔF/F_0_ events for soma (black) and dendrites (orange) for all dendritically amplified (left panel) and dendritically attenuated (right panel) events in a single neuron. ΔF/F_0_ traces are aligned to somatic peak time. Event latency is defined as the time between the somatic and dendritic peaks. Compared to dendritically attenuated events, dendritically-amplified events peaked earlier. **M,** Pearson correlation value between the SD residual and the event latency between soma and its corresponding dendrite indicating that the larger the SD residual, the earlier the dendritic peak is compared to the somatic one (paired t-test, p = 8e^-13^. Mean =-0.075 and - 0.005; SEM = 0.007 and 0.006. n = 466 neurons).

Empirically, we observed that the relative magnitude of coincident events in somas and dendrites varied dramatically, despite event timing correlation being very high (Fig. 2b; consistent with prior studies^39,40,42,45,46,48^). Since event magnitudes at soma and dendrites were best described by a linear relationship (Extended Data Fig. 9 and Extended Data Fig. 10b), we assessed the relative degree of dendritic amplification versus attenuation with a best-fit line through all events and then calculated the somato-dendritic (SD) residual associated with individual transients (Fig. 2b, c)^42^. This captured the variance of dendritic responses for a given somatic event magnitude. We then defined positive and negative residuals as dendritically amplified and attenuated events, respectively.

To test whether SD residuals contain information that is biologically meaningful, we used activity from all the somas in our field of view in the two seconds preceding individual GCaMP events in a neuron of interest (P+ and P- neurons on days 1 to 14) to predict whether these events were dendritically amplified or attenuated (Fig. 2d). To do so, we used a linear Support Vector Machine (SVM), a common algorithm to both classify and regress using high-dimensional data. We found that the performance of our binary classifier on individual neurons strongly correlated with the decoder’s ability to capture the magnitude of dendritic amplification/attenuation in the classification confidence (Fig. 2e, g, h, Extended Data Fig. 10c, d, and Extended Data Fig. 11a, b). This was an emergent property since the decoder was trained for binary classification only and had no information about the magnitude of dendritic amplification/attenuation. Among 466 neurons, approximately 20% showed a significant correlation between classification confidence and the magnitude of SD residual (Fig. 2h, Extended Data Fig. 10c, d, and Extended Data Fig. 11a, b). We found that in these neurons, we could accurately decode 61% of the events as being either amplified or attenuated, well above the 50% chance level (Fig. 2j, Extended Data Fig. 10e, and Extended Data Fig. 11c). Additionally, at the single-cell level, we found a statistically significant positive Pearson correlation between classification confidence and SD residual, demonstrating that the surrounding network of neurons can be used to predict the amplitude of the residual for coincident somato-dendritic transients (Fig. 2k, Extended Data Fig. 10f, and Extended Data Fig. 11d). Importantly, our analysis approach completely decorrelates somatic event magnitude from SD residuals (Fig. 2f, I, and Extended Data Fig. 10a), indicating that mismatches in somato-dendritic coupling are predicted independently from somatic activity and represent information encoded *de novo* in the dendrites. Additionally, our results demonstrate that P_0_ neurons could be decoded at the same level as P+ and P- neurons (Extended Data Fig. 12), and that decoding does not depend on somatic responses to visual stimuli across the three subpopulations (Extended Data Fig. 13).

We further found that dendritically amplified events consistently peaked earlier than dendritically attenuated events compared to the soma (Fig. 2l, m, Extended Data Fig. 10g, and Extended Data Fig. 12e), congruent with results in brain slices^40^.

## Dendritic activity is preferentially reduced by both anesthesia and optogenetic activation of NDNF-positive layer 1 interneurons

Previous studies indicate that anesthesia reduces top-down input and/or inhibits apical tuft dendrites in L5 PNs^22,52–54^. We therefore hypothesized that SD residual should be reduced during anesthesia compared to wakefulness. To test this, we simultaneously recorded somatic and dendritic activity of L5 PNs in RSC during these two conditions (Fig. 3a, b, c). Consistent with previous findings^22^, we observed a dramatic effect of anesthesia on the frequency of GCaMP transients (Fig. 3d). For each neuron, we used all events detected during wakefulness to establish the distribution of SD residuals during awake periods. We then measured the effect of anesthesia on the SD residual using the best-fit somato-dendritic line calculated during wakefulness. Anesthesia strongly reduced the SD residual (Fig. 3c, e), consistent with previous observations of decreased top-down input^22,54^.

**Figure 3:**
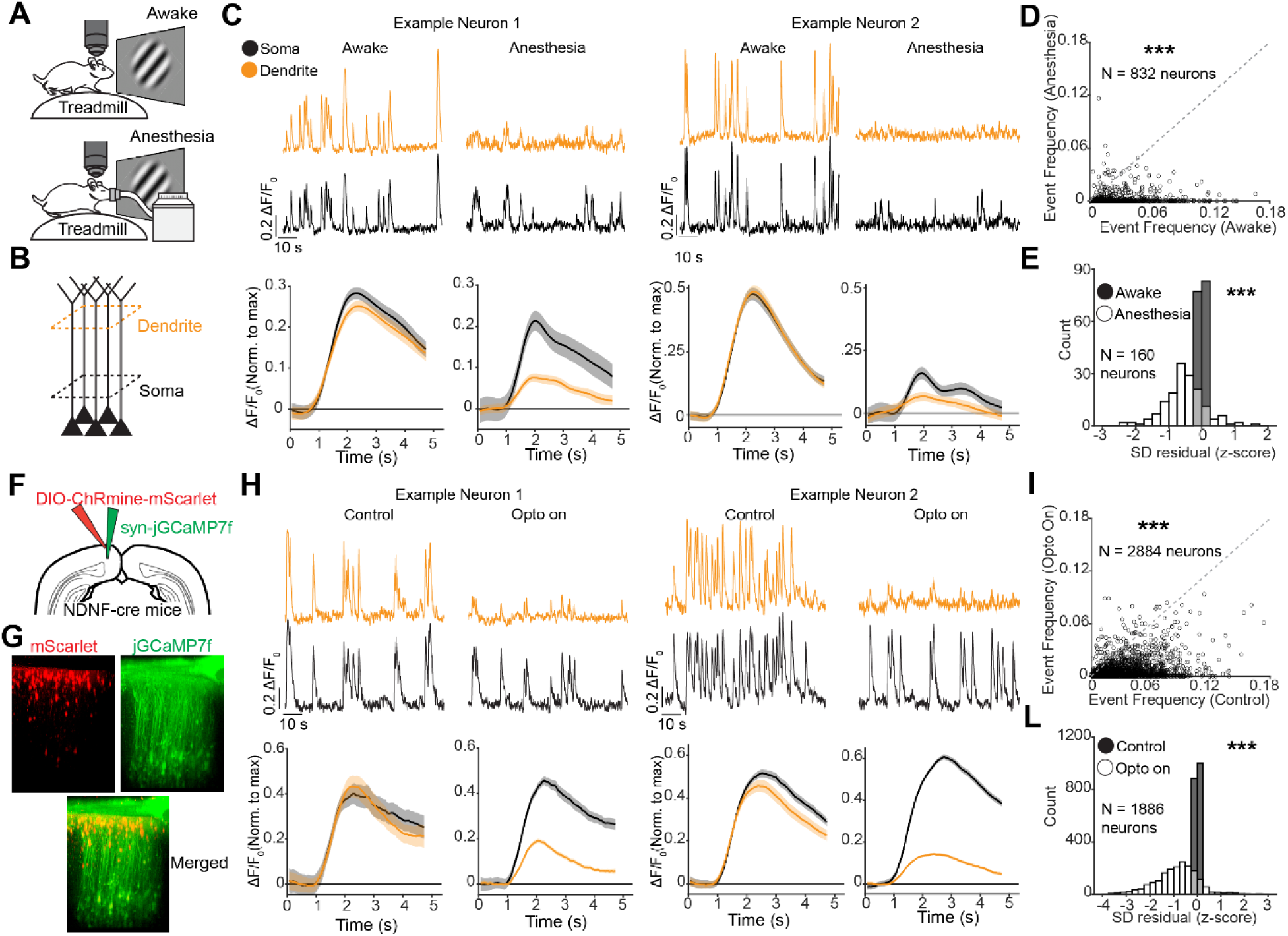
Experimental manipulation of SD residuals. **a,** Schematic of the experimental approach. First, we imaged neurons in RSC while the mouse was exposed to rotating stimuli identical to those presented during baseline estimation in the BCI task. Next, we anesthetized the mice (using isoflurane) and imaged the same neurons. **b,** Schematic of the imaging approach. **c,** Upper panels, ΔF/F_0_ GCaMP traces simultaneously recorded in the soma (black) and its corresponding dendrite (orange) for two representative neurons during wakefulness and anesthesia. Lower panels, the mean ΔF/F_0_ signal in the somas (black) and in the dendrites (orange) of two example neurons, for all events that occurred during wakefulness and anesthesia. Compared to somatic activity, dendrites are preferentially inhibited during anesthesia. Shading represents SEM. **d,** Mean somatic event rate (GCaMP7f) during wakefulness and anesthesia. Dashed grey line represents the identity line. Paired t-test, p < 1.9e^-126^. Mean = 0.002 and 0.0002; SEM = 6.3e^-5^ and 1.8e^-5^. n = 832 neurons for wakefulness and anesthesia, respectively). **e,** Distribution of SD residual during wakefulness (in black) and anesthesia (in white). Paired t-test, p = 3.3e^-20^. Mean = 0 and-0.52; SEM = 2.2e^-18^ and 4.9e^-2^, n = 160 neurons for wakefulness and anesthesia, respectively. **f,** Targeting strategy schematic. A Cre-dependent version of ChRmine tagged with mScarlett was injected into layer 1 of the RSC of NDNF-Cre mice. GCaMP7f under the control of the synapsin promoter was injected into layer 5 of the RSC in the same animals. **g,** Z-stack reconstruction of an imaged field of view. The image was acquired in vivo using 2-photon microscopy (1000 nm laser wavelength). In red, mScarlett. In green, GCaMP7f. **h,** Same as c, for opto on and control conditions. Compared to somatic activity, dendrites are preferentially inhibited during opto on. **i,** Mean somatic event rate (GCaMP7f) during opto on and off. Dashed grey line represents the identity line (paired t-test, p < 2.2e^-308^. Mean = 2.8e^-2^ and 5.9e^-3^; SEM = 4.5e^-4^ and 2e^-4^. n = 2884 neurons for control and opto on, respectively). **l,** Distribution of SD residual during control (in black) and opto on (in white). paired t-test, p = 9e^-214^. n = 1886 neurons, mean = 0 and-0.87; SEM = 7.7e^-19^ and 2.4e^-2^, for wakefulness and anesthesia, respectively.

Prior work has also demonstrated that NDNF-positive layer 1 inhibitory interneurons (L1 INs) can inhibit the apical dendrites of pyramidal neurons^52,53^. We therefore tested whether NDNF- mediated inhibition reduced SD residuals, indicative of a preferential effect on apical dendritic activity. To do so, we co-injected NDNF-Cre mice with both a Cre-dependent version of ChRmine in layer 1 and GCaMP7f, expressed under the control of the synapsin promoter in layer 5 (Fig. 3f, g). We then recorded somatic and dendritic GCaMP activity of individual layer 5 neurons, in the presence and absence of L1 NDNF+ IN activation via an LED light (Fig. 3h). Similar to our approach during anesthesia, we first established the control relationship between somatic and dendritic event amplitudes for each neuron and then compared this to the SD residuals of activity during optogenetic activation. NDNF+ IN activation reduced the frequency of GCaMP transients and, consistent with a number of previous *ex vivo* and *in vivo* studies^52,53,55^, strongly decreased the SD residual in individual layer 5 pyramidal neurons (Fig. 3h, l). Together, these results demonstrate that SD residual is predictably affected by two independent experimental manipulations in vivo, establishing it as a robust metric of dendritic versus somatic activity.

## Somato-dendritic residuals decode reward and trial outcome at the population level

Next, we evaluated if the SD residual contained information about task-related variables that could serve as putative teaching signals. We first tested whether changes in SD residual at the population level contained reward-related information. For each imaging session, we decoded rewarded versus unrewarded trials by comparing the 2 s following neural activity reaching target activation on rewarded trials with the analogous 2 s timeout period during unrewarded trials (Fig. 4a, b, c). Using a linear SVM trained on SD residuals (see Methods), we were able to decode at 63% accuracy on average, above both chance and shuffle performance (Fig. 4d, e, Extended Data Fig. 14, and Extended Data Fig. 15a, b).

**Figure 4:**
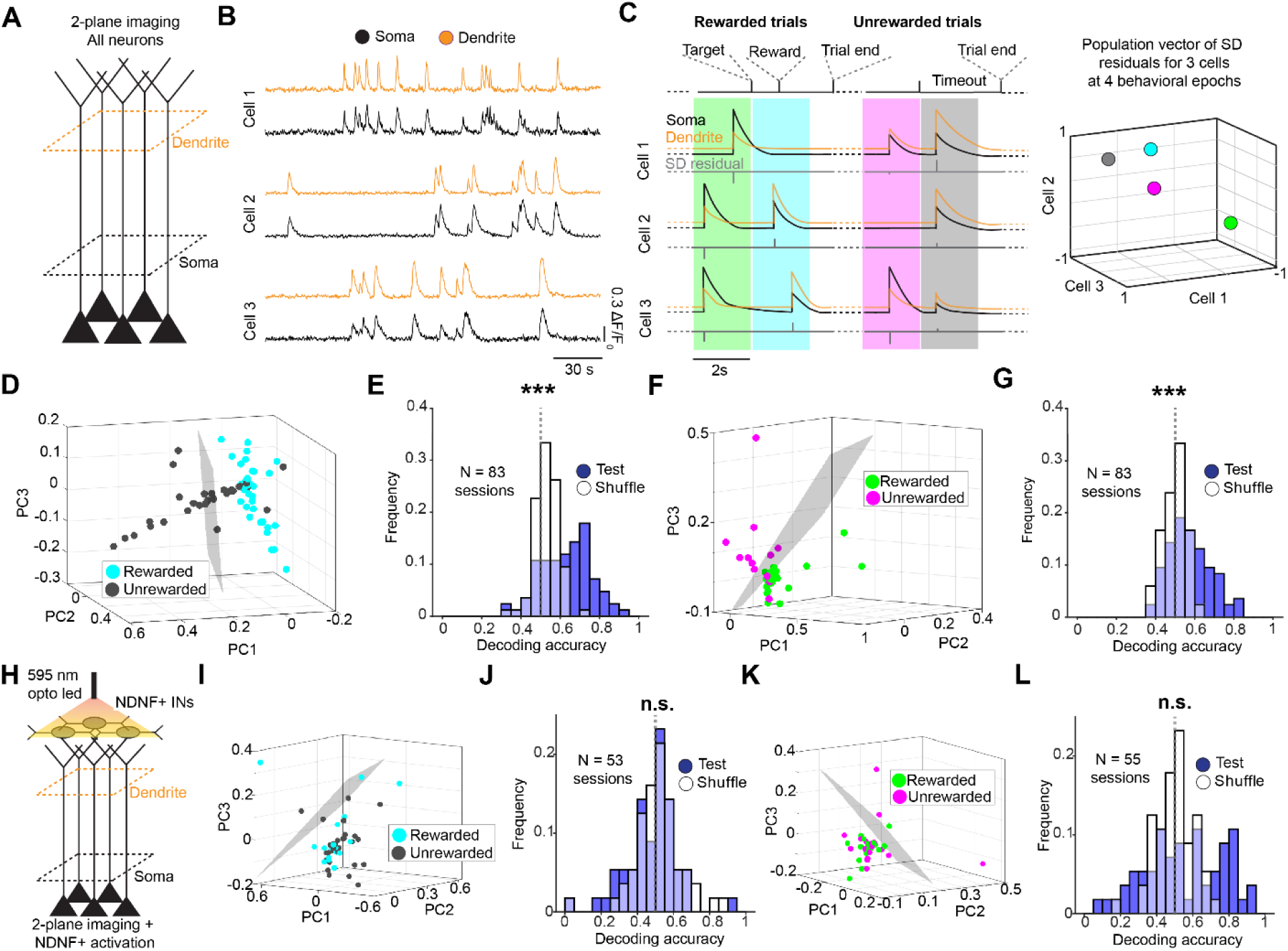
A population vector of somato-dendritic residuals contains reward-related information. **a,** Schematic of the experimental approach. We isolated all neurons in the field of view with a paired soma and dendrite and used the SD residual population vector to decode task-relevant variables. **b,** ΔF/F_0_ GCaMP traces simultaneously recorded in the soma (black) and its corresponding dendrite (orange) for three representative neurons. **c,** Schematic of SD residual population vector for 4 different behavioral epochs. Left panel, in black, orange, and grey the somatic, dendritic, and residual traces for 3 cartoon neurons. The green, cyan, purple and black boxes represent 4 different behavioral epochs: 2 seconds before and after target is reached for rewarded trials and 2 seconds before and after the end of unrewarded trials. Each neuron’s SD residual trace was estimated by collapsing coincident soma-dendrites events into point values at the time of event onset. Resultant SD residual traces for all neurons in each behavioral epoch were used to estimate the n-dimensional vector of SD residuals where n corresponded to the number of neurons for with paired somas and dendrites (see method and figure 2). Right panel, 3-D plot of SD residuals for the four behavioral epochs shown in the left panel, where x, y and z correspond to SD residual of neurons 1-3 from left. **d,** A 39-dimensional vector of SD residuals collapsed onto the first 3 principal components for visualization purposes only. Each dot corresponds to a vector of SD residuals. In cyan, vectors resulting from the two seconds following the reaching of target activity in rewarded trials. In dark grey, vectors calculated in the 2 seconds following the end of an unsuccessful trial (same as C). Shaded black hyperplane represents decision boundary for binary classification calculated using a linear SVM. **e,** Decoding accuracy for test vs. shuffle data for 83 sessions (paired t-test, p = 9.8e^-9^. Mean = 0.63 and 0.52; SEM = 0.01 and 0.01; n = 83 for test and shuffled data, respectively. **f, g,** Same as in d and e: a 264- dimensional vector of SD residual collapsed onto the first 3 principal components for visualization only. Green represents the last two seconds of a rewarded trial while purple represents the last two seconds of an unrewarded trial (paired t-test, p = 7.1e^-8^. Mean = 0.57 and 0.49; SEM = 0.01 and 0.01; n = 83 for test and shuffled data, respectively). **h,** Schematic of the experimental approach: L1 NDNF+ INs were optogenetically activated during BCI task performance. **i, j, k, l,** Same as d-g but during optogenetic activation of NDNF+ INs. In j, paired t-test, p = 0.18. Mean = 0.48 and 0.51; SEM = 0.02 and 0.02; n = 53 sessions for test and shuffled data, respectively. In l, paired t-test, p = 0.13. Mean = 0.54 and 0.50; SEM = 0.03 and 0.01; n = 55 sessions for test and shuffled data, respectively.

Next, we tested if inputs onto the apical tuft dendrites represent instructive signals during learning. We used SD residuals to decode successful versus unsuccessful trials in the 2 s periods preceding successful target activation versus timeout, respectively. Once again, we found that our decoder performed significantly above chance at 57% accuracy on average (Fig. 4c, f, g, Extended Data Fig. 14 and Extended Data Fig. 15c, d), demonstrating that individual neurons encode information about the network states that correspond to successful versus unsuccessful outcomes in their SD residuals both before and after reward delivery. Since the trial time we analyzed is pre-outcome, our results indicate that the SD residuals encode instructive signals based on the task-associated reward function.

Finally, we tested the role of L1 inhibition in controlling dendritic signals encoding reward and trial outcome (Fig. 4h). To do this, we performed experiments on a second set of mice expressing ChRmine in NDNF+ L1 INs. Optogenetic activation of L1 NDNF+ neurons abolished task and reward-related information in the apical dendrites of layer 5 pyramidal neurons (Fig 4i, j, k, l), indicating a key role for local cortical inhibition in dendritic processing task-related variables.

## Somato-dendritic residuals reflect neuron-specific task error signals

We exploited the explicit definition of error and of functionally-opposite classes of neurons in our experimental design to test whether error signals are received at apical dendrites and, if so, whether they differ between neurons according to each neuron’s causal role in the task (Fig. 5a, b). We reasoned that a scalar error signal would manifest as amplified dendritic activity during periods of error reduction for both P+ and P- neurons and as attenuated dendritic activity during times of error increase. However, a vectorized error signal would exhibit selective P+ versus P- dendritic activation, since the activity of each group is causally mapped to error in opposite ways. To disambiguate between these scenarios, we averaged the error in 2 s windows throughout the task and defined each window as an error increase or decrease epoch, given that the angle of the visual stimuli presented to the animals represented the instantaneous task-associated error (Fig. 5a). Next, we calculated the SD residuals for P+ and P- neurons for coincident soma-dendrite events in each window during error decrease and error increase epochs. Since our analysis was restricted to time bins with coincident somato-dendritic events in P+ and/or P- neurons, any potential noise-driven flickering was not present in our analysis. We found that the dendrites of P+ neurons were relatively amplified during error-reduction compared to error-increase epochs (Fig. 5c). Dendrites in P- neurons exhibited the converse relationship: relative dendritic attenuation and amplification occurred during error-reduction and error increase, respectively (Fig. 5d, e, f, Extended Data Fig. 16, and Extended Data Fig. 17). This relationship could be observed in 6 out of 6 mice trained in the task (Extended Data Fig. 18) and remained intact when we restricted our analysis to neurons whose somatic activity was the same during epochs of error increase and reduction (Extended Data Fig. 19). Additionally, the same inverted relationship between dendritic signals and task-associated errors was found in the dendrites of P_0_ neurons which were functionally correlated to P+ and P- neurons (Extended Data Fig. 20). Intriguingly, SD residuals represented error derivatives, not errors (Extended Data Fig. 21), in contrast to instructive signals found in the classical implementations of backpropagation.

**Figure 5:**
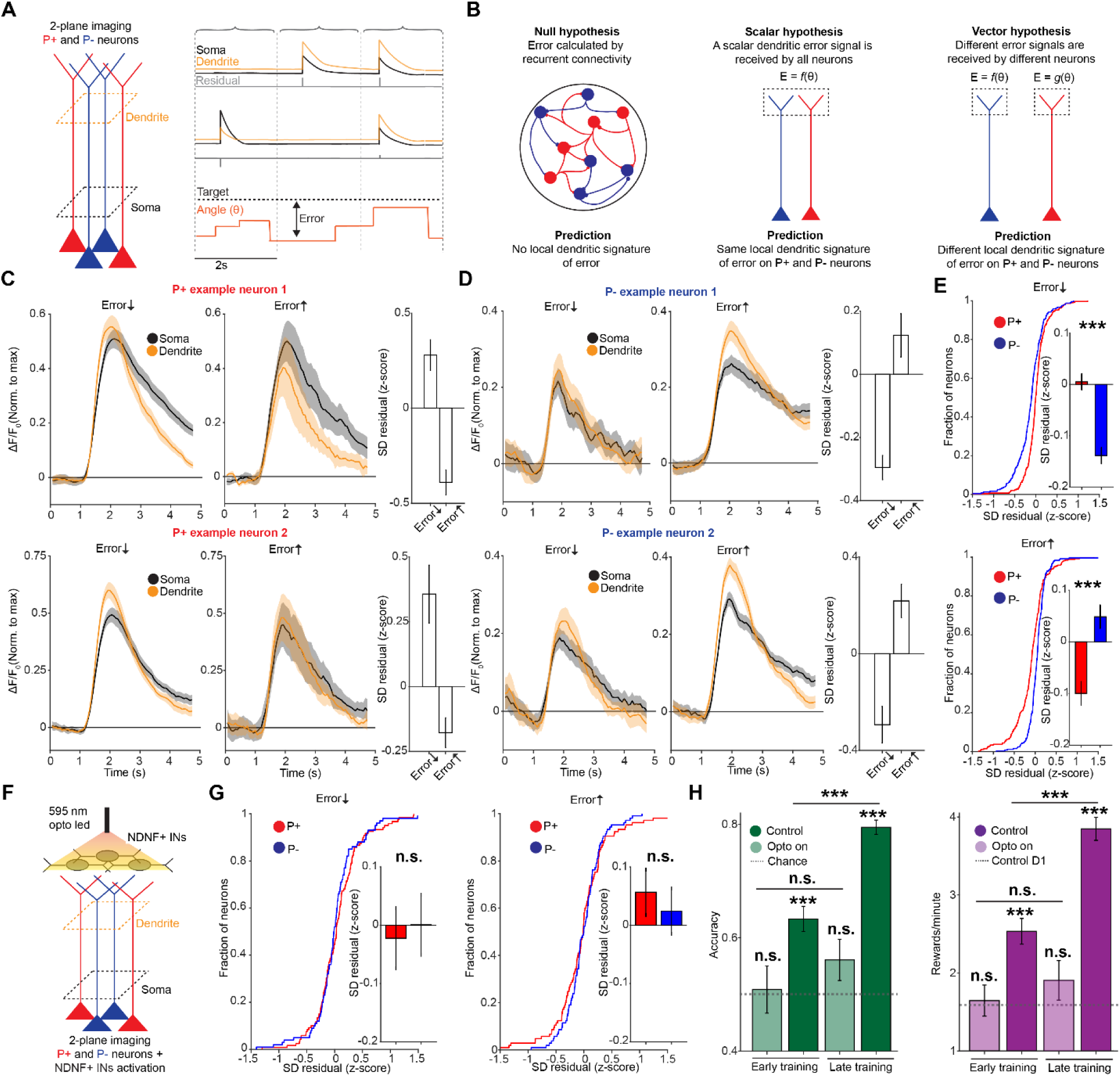
Dendritic error signals are cell-specific and depend on the causal contribution of the neuron to the task. **a,** Experimental schema. Black, orange and grey indicate idealized somatic, dendritic and residual traces for 3 neurons. Relationship between stimulus angle, target, and error shown below. All closed-loop data from the BCI paradigm (which excludes rewards and timeout periods) were chunked into 2 seconds bins. Epochs of error decrease and increase were defined as bins in which the mean derivative of the angle increased and decreased, respectively. **b,** Three possible hypotheses: In the null hypothesis scenario, error is calculated at the population level through recurrent dynamics independent of dendrites. In the scalar hypothesis, a single error signal is broadcasted through the dendrites of all neurons in the network. This model predicts that all neurons receive the same error signal. In the vector hypothesis, error signals received on the dendrites of individual neurons are tailored according to the causal involvement of individual neuron to behavior. This model predicts that neurons with opposite causal contribution to behavior will receive different error signals onto their dendrites. **c,** For two individual P+ neurons, the mean ΔF/F_0_ signal in the somas (black) and in the dendrites (orange) for all events that occurred during epochs of error reduction and error increase respectively. Compared to somatic activity, dendrites are relatively amplified during error reduction compared to error increase epochs. The bar graph represents the mean SD residual value (z-scored) for all events occurred during error decrease and increase epochs. Error bars represent SEM. **d,** Same as c for two P- neurons. Contrary to the P+ neurons, dendritic activity is relatively attenuated for error reduction epochs compared to error increase epochs. **e,** Left panel, during error reduction epochs, the cumulative distribution function for SD residuals (z-scored across all neurons) for P+ (in red) and P- (in blue) neurons. The bar graph represents the mean and SEM for the population distribution shown in the cumulative distribution function for P+ and P- neurons. Dendrites of P+ neurons are relatively amplified compared to the dendrites of P- neurons during error reduction epochs (t-test; p = 5.3e^- 7^; mean = 0.005 and-0.14; SEM = 0.01 and 0.02; n = 292 and 240 for P+ and P- neurons respectively). Right panel, error increase epochs. Dendrites in P+ neurons are more attenuated than P- neurons during error reduction epochs (t-test, p = 1.2e^-7^; mean =- 0.1 and 0.05; SEM = 0.02 and 0.01; n = 267 and 249 for P+ and P- neurons respectively). **f,** Experimental schematic: GCaMP signals in the soma and dendrites of P+ and P- neurons were recorded during optogenetic activation of L1 NDNF+ neurons. **g,** As in e but during L1 NDNF+ neuron activation: left panel, during error reduction epochs, the cumulative distribution function for SD residuals (z-scored across all neurons) for P+ (in red) and P- (in blue) neurons. The bar graph represents the mean and SEM for the population distribution shown in the cumulative distribution function for P+ and P- neurons (t-test; p = 0.58; mean = 0.06 and 0.02; SEM = 0.04 and 0.04; n = 119 and 100 for P+ and P- neurons respectively). Right panel, same as e, for error increase epochs. Dendrites in P+ neurons are more attenuated than P- neurons during error reduction epochs (t-test, p = 0.7; mean =- 0.02 and 0; SEM = 0.05 and 0.03; n = 105 and 105 for P+ and P- neurons respectively). **h,** BCI performance. Left panel, (accuracy, in green) in early and late training for control (opto off) and opto on condition. Dashed grey line represents chance level (paired t-test; p = 0.83; mean = 0.51, SEM = 0.04; n = 32 for opto on against chance during early training; paired t-test; p = 3e^-7^; mean = 0.63, SEM = 0.03; n = 48 for control against chance during early training; p = 0.1; mean = 0.56, SEM = 0.02; n = 24 for opto on against chance during late training; p = 5.7e^-23^; mean = 0.8, SEM = 0.01; n = 36 for control against chance during late training; t-test, p = 0.36 and 1.3e^-7^ for early vs late training during opto on and control condition, respectively). Right panel, rewards per minute (in purple) in early and late training for control (opto off) and opto on condition. Dashed grey line represents rewards per minute for control on day 1 (paired t-test; p = 0.8; mean = 1.6, SEM = 0.2; n = 32 for opto on against control on day 1, during early training; paired t-test; p = 7e^- 7^; mean = 0.63, SEM = 0.26; n = 48 for control against control on day 1, during early training; p = 0.23; mean = 1.9, SEM = 0.26; n = 24 for opto on against control on day 1, during late training; p = 4.2e^-17^; mean = 3.8, SEM = 0.14; n = 36 for control against control on day 1, during late training; t-test, p = 0.42 and 1.5e^-7^ for early vs late training during opto on and control condition, respectively).

Next, we tested whether vectorized error-related dendritic signals were necessary for learning by optogenetically activating NDNF+ L1 INs throughout the BCI task. This abolished vectorized error-related signaling in the apical tuft of layer 5 PNs (Fig. 5f, g) and disrupted learning (Fig. 5h). This demonstrates that local computation in the apical dendritic tuft is necessary for performance improvements in the BCI task.

## Discussion

Here, we demonstrate the first use of neurofeedback brain computer interfaces to study the mechanisms of biological credit assignment at the subcellular level. Our results provide the first biological evidence of a vectorized solution to the credit assignment problem in the brain via cortical dendrites. Our data is consistent with a model of credit assignment in which learning is instructed by instantaneous, vectorized teaching signals received onto the distal dendrites of pyramidal neurons^1–5^. This spatial segregation mechanism allows cortical circuits to overcome the biologically implausible temporal separation of feedforward and feedback streams conventionally used for computing teaching signals during vectorized learning in ANNs.

The data presented here reveal magnitude differences in coincident somato-dendritic events that can be predicted using activity in the surrounding network of neurons. At the population level, differences in somato-dendritic coupling encode *de novo* information relative to somatic activity. This information could be used by individual neurons as instructive signals, such as reward and task error, providing novel evidence that individual neurons can explicitly access the reward function of a learning task through independent dendritic computation. We further demonstrate that cell-specific changes in SD residuals correlate with the functional role of individual neurons as well as with subsequent changes in activity levels during learning. Finally, optogenetic activation of NDNF+ L1 INs disrupted both dendritic computation and learning, demonstrating that dendritic processing is necessary for learning.

Our results demonstrate the existence of a signed, vectorized dendritic input that is tailored in a condition-specific manner to individual neurons. The extent to which this dendritic activity reflects moment-to-moment computational signals – as opposed to teaching signals for synaptic weight changes – remains to be elucidated. Future work is needed to assess whether these dendritic signals result from glutamatergic inputs from higher-order cortical areas, from neuromodulation, or as a product of recurrent excitatory and inhibitory local computation. Dopaminergic signaling specifically has been causally implicated in both error signaling and in learning neurofeedback BCI tasks in rodents and humans^32,56–58^ and thus represents a compelling target for future investigation. Further experiments are also needed to test whether errors signals are calculated locally at each hierarchical layer or are transmitted across layers, as in the classical formulation of backpropagation^19^. Previous neurofeedback BCI studies have demonstrated that degrading the contingency between neuronal activity and feedback stimuli impairs learning^32–34^: future work will have to determine whether external stimuli are always necessary for error representations or whether animals can access the cost function via internal states exclusively, and how the dendritic representation of error might change as a result.

The error signals we observed have appealing connections to the gradient calculations found in the backpropagation algorithm. In contrast to the classical implementation of backprop, however, we observed that dendrites received signals that bore signatures of error derivative rather than error itself. Intriguingly, our results could also be consistent with target propagation (specifically, difference target propagation)^14,20,21^. Indeed, our data indicate that dendritic activity contains a target signal for the parent soma in addition to task-related error information. Future approaches, built on the framework we present here, could be used to disentangle the specific learning algorithm(s) employed by the brain^14,59^.

Together, our results help to reconcile early findings and theories of dendritic function, which focused on single dendritic branches as the building blocks for independent computation, with later in vivo findings that have demonstrated prevalent co-occurrence of dendritic and somatic events^15,24,50,60^. By demonstrating that apical dendrites locally compute reward and error-related signals, our results present a framework for dendritic computation which does not require fully independent dendrites to perform credit assignment for adaptive behavior and highlight new directions for the development of biologically-inspired ANNs.

## Methods

### Animals

All experiments were compliant with guidance and regulation from the NIH and the Massachusetts Institute of Technology Committee on Animal Care. Male and female Rbp4-Cre and NDNF-Cre heterozygous mice were maintained on a 12-hour light/dark cycle in a temperature-and humidity-controlled room with ad libitum food access and were used for experiments at 8-15 weeks of age. Except for anesthesia experiments, animals were water-deprived by decreasing water intake from 3 ml to 1.2 ml over the course of 10-14 days and maintained at 1.2 ml thereafter, for 5-7 days before experiments and throughout training.

### Surgery

Mice were initially anaesthetized using 4% isoflurane and subsequently maintained at 1-2% isoflurane through the rest of the surgery. Body temperature was maintained at physiological levels using a closed-loop heating pad. Additional heating was provided for post-surgical recovery. To protect eyes from dryness, eye cream (Bepanthen, Bayer) was applied. Animals were injected with Dexamethasone (4mg/kg), Carprophen (5mg/kg) and Buprenorphine (slow release, 0.5mg/kg) subcutaneously. The scalp was shaved using hair-removal cream and cleaned afterwards using iodine solution and ethanol. Next, the skull was exposed. For in vivo imaging, a 3mm-wide craniotomy was performed. In Rbp4-Cre mice, at 3-4 different sites, we injected 100 nl of AVV1-syn-FLEX-jGCaMP7f-WPRE (Addgene, catalog # 104492-AAV1, 2-5×10^12^ vg/ml concentration after a 1:10 dilution from the original concentration) at 400 μm from the surface of the brain in the left hemisphere of the Retrosplenial cortex (2.5 mm caudal of bregma). The same labeling approach was utilized to perform anesthesia experiments. In NDNF-Cre mice, we injected 100nl of AAV8-nEF-Con/Foff 2.0-ChRmine-oScarlet (Addgene, catalog # 137161-AAV8, 7×10^12^ vg/ml after a 1:5 dilution from the original concentration) 150 μm from the surface of the brain and 75nl of AAV1-syn-jGCaMP7f-WPRE (Addgene, catalog # 104488-AAV1, 2-5×10^12^ vg/ml concentration after a 1:10 dilution from the original concentration) 500 μm from the surface of the brain. The dura was left intact. Cranial windows consisted of two stacked 3mm coverslips (inserted within the craniotomy) attached to a larger 5 mm coverslip which was subsequently fixed to the skull using cyanoacrylate glue and dental cement. A custom metal headplate was implanted in order to perform imaging under head-fixed conditions. At the end of the procedure, a single dose of 25mL/kg of Ringer’s solution was injected subcutaneously to rehydrate the animal. Recordings started 4-6 weeks post-surgery.

### Two photon imaging

A Neurolabware 2-photon microscope equipped with GaAsP photomultiplier tubes was used for data acquisition. Imaging was performed at 980 nm using an ultrafast pulsed laser (Spectra-Physics, Insight DeepSee) coupled to a 4x pulse splitter to reduce photodamage and bleaching. For excitation and photon collection we used a 16x Nikon objective with 0.8 numerical aperture. Bidirectional scanning was performed (512×796 pixels) semi-simultaneously in two separate planes using an electrically-tunable lens at 30.92 Hz (15.46 Hz for each plane). Laser intensity was independently optimized at each imaging plane using an electro-optical modulator. A custom light shield was attached to the headplate in order to avoid light contamination. Animals were habituated to human handling for 5-10 minutes every day and to head-fixation for 15 minutes a day for at least 3 days directly preceding imaging. Small 10% sucrose water rewards were randomly dispensed during habituation. Daily water intake of at least 1.2 mL was maintained throughout the behavioral experiments. The animal’s locomotion was recorded using an optical encoder (E6, US Digital, 2500 cycles per revolution) tracking the rotation of a cylindrical treadmill 19 cm wide in radius and acquired using the Scanbox software interfaced to a custom-built Arduino system. To maximize the number of units recorded while simultaneously reducing signal contamination, we imaged the trunk of layer 5 pyramidal neurons at two different planes: proximal to the soma and right below the nexus (tuft bifurcation point).

### Optogenetic stimulation

For optogenetic stimulation we employed a Cyclops LED driver from open ephys (Catalog # OEPS-6602) triggered using a direct 6ms TTL pulse delivered via the Neurolabware Dual PSOC box. The driver controlled a fiber-coupled 595 nm LED laser (8.7mW, 100mA ThorLabs catalog # M595F2). LED activation was synchronized with the PMTs of the imaging system using custom-made Matlab scripts. In brief, for every new frame acquired by the 2-photon microscope, the LED was activated for the initial 6ms of the frame, while the PMTs were kept shut off for 1 additional millisecond (7ms total of PMT off time). PMTs would then reactivate to collect calcium data for the remaining ∼24ms of the ∼31ms frame.

### Brain computer interface task

Similar to previous implementations of brain-computer interface learning paradigms^33,34,37^, mice were trained so that they obtained rewards by modulating the activity of 8 or 10 layer 5 pyramidal neurons in the retrosplenial cortex to control the rotation of a grating Gabor patch. The 8 or 10 neurons were equally divided into 2 subpopulations, P+ neurons whose activity rotated the stimulus towards a target angle of 90-degrees (horizontal) and P-neurons whose activity rotated the stimulus away from the target angle, towards a 0-degree (vertical) orientation. Neural activity was transformed into a visual stimulus angle according to the following method: At the beginning of each session, we measured the baseline responses of P+ and P- neurons to 7 randomly presented oriented gratings (0-, 15-, 30-, 45-, 60-, 75-, 90-degree, passive viewing) for approximately 13 minutes (12000 frames). ΔF/F_0_ was calculated for individual P+ and P- neurons and averaged across each population. The mean P- population signal was subtracted from the mean P+ population signal. Next, we randomly resampled 200 trials (435 frames each) from the aforementioned 12000-frame baseline recording and iteratively searched (in 0.005 ΔF/F_0_ incremental steps) for the subtracted ΔF/F_0_ value producing a 50% success rate. That value was set as the threshold value for target activity during the closed-loop phase of the BCI task. Next, we calculated the mean and standard deviation of the subtracted ΔF/F_0_ signal distribution and created a new distribution by mirroring the left side to the right. On day 1, we estimated the z-score corresponding to the ΔF/F_0_ threshold value on the mirrored distribution. On the following days, we estimated the subtracted ΔF/F_0_ signal distribution and its corresponding left-mirrored distribution in the same way as described above in the same way as described above, and utilized the ΔF/F_0_ value corresponding to the z-score used on day 1 as the task target activity during the closed-loop phase of the task. In this way mice could learn the task by either decreasing activity of P- neurons or increasing activity in P+ neurons (or both). The mapping between neuronal activity and visual feedback angle was defined as follows: 0-degree angle corresponded to the minimum value in the subtracted ΔF/F_0_ signal distribution while target, or 90-degree angle was reached at subtracted ΔF/F_0_ value corresponding to threshold defined as described above. Activity in between was split into 7 equally spaced bins each corresponding to a 15-degree interval between 0 and 90 degrees. At each screen refresh, the angle presented reflected the mapping between angle bins and the subtracted ΔF/F_0_ signal averaged over the last 3 frames. The screen refreshed every time a 2-photon frame at the soma was acquired (at 15Hz). In line with previous studies performing neurofeedback BCI in rodents^32–35,37^, we binned the visual stimulus to avoid noise-driven, frame-by-frame stimulus updates at the screen refresh rate, which is beyond the perceptual threshold in mice^61,62^. To avoid introducing a second, orthogonal dimension to our task that would disrupt the straightforward mapping between neuronal activity and task error, we did not introduce any requirement on the number of P+ or P- neurons required to be simultaneously active to trigger a reward or a stimulus update. In each trial, mice had 28 seconds to reach target activity. If they did, a reward, consisting of 4 μL of 10% sucrose water was delivered 1 second after. Additionally, after reaching target activity, the stimulus froze to a 90-degree angle for 2 seconds. After that, mice saw a black screen for 1 additional second and a new trial was initiated. All new trials were initiated by a 0.5 s iso-luminant grey stimulus. If a mouse did not reach target activity within the 28 seconds of the trial, a 3 seconds timeout was given to them consisting of a black screen. To avoid the problem of drifting baselines, ΔF/F_0_ for each neuron was calculated as (F_i_ – F_i_0)/F_i_0 where F_i_0 was the 10^th^ percentile of fluorescence in the previous 30 seconds. For the optogenetics experiments, we recorded 2 different baselines (during passive viewing). The first one with the LED off (control) was used for post-hoc analysis only. The second baseline was recorded during LED stimulation (opto on) and was used to map neuronal activity to angle and target during the closed-loop part of the BCI task. The closed-loop part of the BCI task was recorded during the opto on condition only. Early and late training were defined as days 1-8 and 9-14 respectively, based on average performance (accuracy) remaining above 0.75 in the control (opto off) condition. For anesthesia experiments, we recorded two passive viewing sessions where we presented the same set of stimuli presented when recording the baseline session for the BCI task. We anesthetized the animals in between these two sessions by initially administrating (via inhalation) 4% isoflurane that subsequently decreased to 1% for the duration of the imaging session.

### P+ and P- neuron selection

On day 1, we drew 20-40 ROIs in a single field of view prior to starting our baseline recording. Next, we recorded a session of passive visual stimuli that we would later use as our baseline recording for day 1. At the end of this recording, all ΔF/F_0_ traces for all drawn ROIs were plotted and visually inspected using a custom Matlab script. The experimenter would then select either 8 or 10 of these traces based on event frequency, SNR (determined as the ratio between noise band width and maximum event size), baseline stability, and calcium transient dynamics (with a clear rise, peak, and exponential decay – as opposed to plateau-looking events). The best 8-10 neurons would then be selected from the available pool of neurons on which ROIs were drawn. No arbitrary parameter cutoff (e.g. minimum event frequency or SNR) was introduced. The subdivision of these 8-10 neurons into the P+ and P- population would then be determined by a random number generator. Once selected, P+ and P- neurons would remain the same for the entire duration of the experiment.

### Online motion-correction

In order to avoid drifts in x and y out of our selected regions of interest, we used a Fast-Fourier transformation approach to live motion-correct our movies. To do so, at the beginning of each recording session we acquired a reference image by averaging 20-40 seconds (300-600 frames) collected onto our field of view. To motion-correct each subsequent frame, we selected 4 smaller central areas to register independently from one another (2-D rigid translation) against the corresponding 4 areas in the reference image^63^. We finally rigid-translate the entire 2-D image by taking the average translation in x and y for these 4 subregions.

### Visual stimuli

Visual stimuli were generated using the Psychophysics Toolbox package for MATLAB (MathWorks, MA)^64^ and displayed on a monitor 20 cm away from the contralateral eye. Visual stimuli consisted of a rotating Gabor patch at 7 angles spaced 15 degrees apart from 0 to 90.

### Offline image analysis and signal extraction

To correct for brain motion after image acquisition, as well as to automatically detect ROIs, we used the Suite2P pipeline^65^. For each field of view (FOV), we removed duplicates by excluding ROIs whose signal correlation was above 0.6 and whose center was within 20 μm of distance. In order to separate trunk signals from potential neuropil contamination, fluorescence signals of our ROIs were processed using FISSA^66^ with the following parameters: 4 neuropil subregions and alpha = 0.1. To estimate ΔF/F_0_ after neuropil subtraction, we calculated ΔF/F_0_ at time point i as (Fi – F0)/F0. F0 is defined as the 10^th^ percentile of a 120 seconds long sliding window to remove fluorescence drifts over the course of imaging. Next, we performed spike inference using the CASCADE model Global_EXC_15Hz_smoothing200ms^44^.

### Field of view matching and ROI registration across days

Registration of neurons across days for BCI training was performed manually at the beginning of each session with the help of a custom-designed software. On day 1, a mean intensity reference image of our field of view of interest was acquired. Using a custom-design software, we manually drew 10-20 reference ROIs which included any recognizable brain structure including dendrites, cell somas and sharp-contrast blood vessels. On the following days, after manually finding the same approximate area for the field of view imaged on day 1, a more accurate manual registration was performed by aligning our reference ROIs drawn on day 1 with their corresponding structures on following days. As the relative x and y distance between structures varies along the z-plane, our approach allowed us to consistently match our field of view on day 1 across x, y and z dimensions on any given day. Offline registration of ROIs across days on the other hand, was initially performed using the ROIMatchPub implementation for Suite2P followed by an exhaustive manual curation.

### Quantification of event frequency, magnitude, and timing

Events were detected for each ROI using the MATLAB function *findpeaks* on the spike-inferred signal. For analysis of the spike inferred signal, we estimated the integral of individual peaks by multiplying the height and width of individual transients. Event occurrence was defined as the time at which spike probability peaked. For ΔF/F_0_ analysis, once we found an event, we utilized a 2-seconds backward sliding window to identify the frame at which the derivative of the ΔF/F_0_ signal became consecutively positive for 300 ms. This was considered the transient onset frame while the peak of the transient was considered the maximum ΔF/F_0_ value in the 2 seconds following peak detection. We therefore estimated the integral of the ΔF/F_0_ signal by multiplying the height (maximum ΔF/F_0_ value – ΔF/F_0_ value at transient onset) and the width (frame at maximum ΔF/F_0_ value – frame at transient onset) of the ΔF/F_0_ signal. The backward and forward detection windows were limited in time by the presence of a precedent or subsequent event detected using the spike-inferred signal. Proximal trunks were paired to their correspondent distal trunk whenever their ΔF/F_0_ signal correlation was equal above 0.6. For optogenetics and anesthesia experiments, we matched proximal trunks to their corresponding distal trunk using activity during control (opto off) and wakefulness, respectively. Whenever we found more than 1 distal dendrite correlated with the same proximal trunk, we selected the one with the best signal-to-noise ratio, so to always have a single distal dendrite associated with a proximal trunk. Coincident events were defined as two events occurring (independently detected) within a 500 ms window in the two compartments. To quantify the somato-dendritic magnitude mismatches of coincident events, we first fit a best-fit line against the somatic and dendritic magnitudes of all events. For each event, we calculated the residual from the best-fit line, and defined residuals larger than 0 as dendritically-amplified and residuals smaller than 0 as dendritically-attenuated. To estimate the SD residual during optogenetic stimulation and anesthesia, first we estimated the best-fit line using somato-dendritic activity during light off and wakefulness conditions, respectively. Next, we calculated the residual for all events detected during opto on and anesthesia as the distance between these events and the previously-calculated best-fit line.

### Decoders

To decode whether individual transients would be amplified or attenuated, we trained a support vector machine binary classifier (SVM, linear kernel) using stochastic gradient descent^67^ (as implemented by MATLAB *fitclinear*). For each coincident event in the soma and dendrites, we averaged the spike-inferred activity of each neuron in our field of view (excluding the neuron of interest) in the 2 preceding seconds, and we used this average activity to create an n-dimensional population activity vector where n corresponds to the number of isolated units in our field of view. The binary classifier was trained to separate dendritic amplification from dendritic attenuation (see above) using a leave-one-out approach. Accuracy was determined as the fraction of correctly classified events. For imbalanced datasets, we used a Synthetic Minority Oversampling Technique (SMOTE, k neighbors = 5) to train (not test) using a balanced dataset. SMOTE was applied after separating our train and test dataset. To control for any potential data leakage, our shuffle control went through the exact same procedure as our test dataset, including SMOTE oversampling with the only difference that labels were randomly shuffled before separating the train and test data. We calculated the confidence of a prediction as the Euclidean distance from the hyperplane. Reward-associated and reward-instructive epochs were defined as 2 seconds before and 2 seconds after the reach of target activity, respectively for successful trials, and 2 seconds before and 2 seconds after the end of a trial for unsuccessful trials. To decode successful from unsuccessful trials, we generated a n-dimensional somato-dendritic residual vector by taking the residual for each neuron for which we identified a somato-dendritic pair (see above) in these two seconds epochs. Neurons inactive in the two seconds epochs were assigned a value of 0. The binary classifier was trained in the same manner as described above.

### Statistics

All analysis was performed using MATLAB 2020a using custom-written scripts and functions. All error bars in figures represent standard error of the mean (SEM). Statistical tests and independent samples are described in figure legends.

## Data and code availability

All analysis, BCI code, and data are available upon request.

## Acknowledgements

We thank Ila Fiete and Courtney Yaeger for comments on the manuscript. V.F. is Y. Eva Tan Molecular Therapeutics Postdoctoral Fellow. V.D.T. is a Janet and Sheldon (1959) Razin Fellow.

E.H.S.T. was supported by the National Institute of General Medical Sciences (T32GM007753) and the Paul & Daisy Soros Fellowship. M.T.H. was supported by the NIH (RO1NS106031, R01NS113079, and R01MH135141) and by the Klingenstein-Simons Fellowship, the Vallee Foundation Scholars, and the McKnight Scholars programs.

## Contributions

V.F. conceptualized and designed the experimental approach, designed the BCI, habituated and performed surgery on mice, collected the data, conceptualized and implemented the analyses, prepared the figures and wrote the manuscript. V.D.T. helped in the conceptualization of data analysis, building the data analysis pipeline and writing the manuscript. N.J.B. performed surgeries on mice. E.H.S.T. helped write the manuscript. M.T.H supervised all aspects of the project.

## Ethics declaration

The authors declare no competing interest.

## Supplemental Materials

**Supplementary Figure 1:**
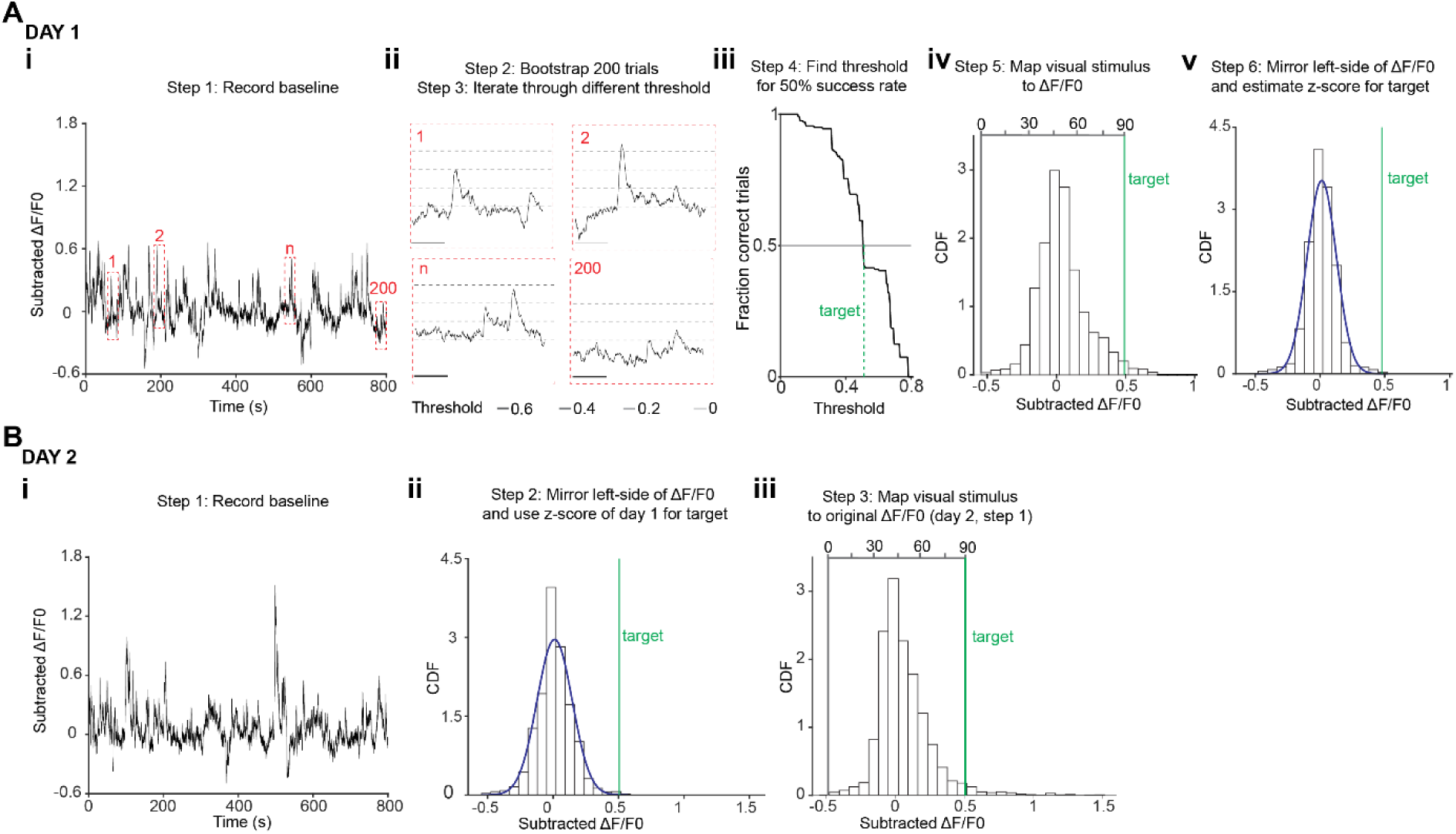
Determining target activity and mapping visual feedback to ΔF/F0 within and across days. a, Mapping strategy on day 1. **ai**, A baseline of approximately 13 minutes (12,000 frames imaged at ∼15Hz) was recorded each day for P+ and P-neurons, and the mean ΔF/F0 for each subpopulation was subtracted from one another (also see Fig. 1e). **aii,** The subtracted signal was used to bootstrap 200 trials (28 seconds each resampling with replacement 200 start times, red boxes). Next, we iteratively searched across different thresholds. **aiii,** For each threshold (in incremental steps of 0.005 ΔF/F0), we selected the one which was crossed 50% of the times in the 200 resampled trials. This value was set as the target value (i.e. 90-degree angle). **aiv**, Next we mapped the subtracted ΔF/F0 to a visual feedback stimulus. 90-degree was mapped as the ΔF/F0 at which the animal reached target on 50% of the resampled trials (see aii), 0-degrees was mapped as the minimum value in the ΔF/F0 distribution. The space in between was linearly spaced into 6 bins whose edges marked the boundary between 15-degrees angles. **av**, The left side of the subtracted ΔF/F0 was mirrored on the right and fitted using a Gaussian function. The z-score of the target value (see aii), was recorded for mapping on subsequent days. **b**, Target activity and mapping strategy on day 2-14. **bi**, We recorded baseline activity in the same way as shown in ai. **bii**, After estimating the distribution of the subtracted ΔF/F0 signal, we mirrored the left part of the distribution and fitted a Gaussian function as described in aiv. Our target was defined as the same z-score of the target value on day 1 (see iv). **biii**, On the original subtracted ΔF/F0 distribution, we mapped 90-degrees as the z-score of the target value on day 1 (see bii) and 0-degrees as the minimum value in the subtracted ΔF/F0 distribution.

**Supplementary Figure 2:**
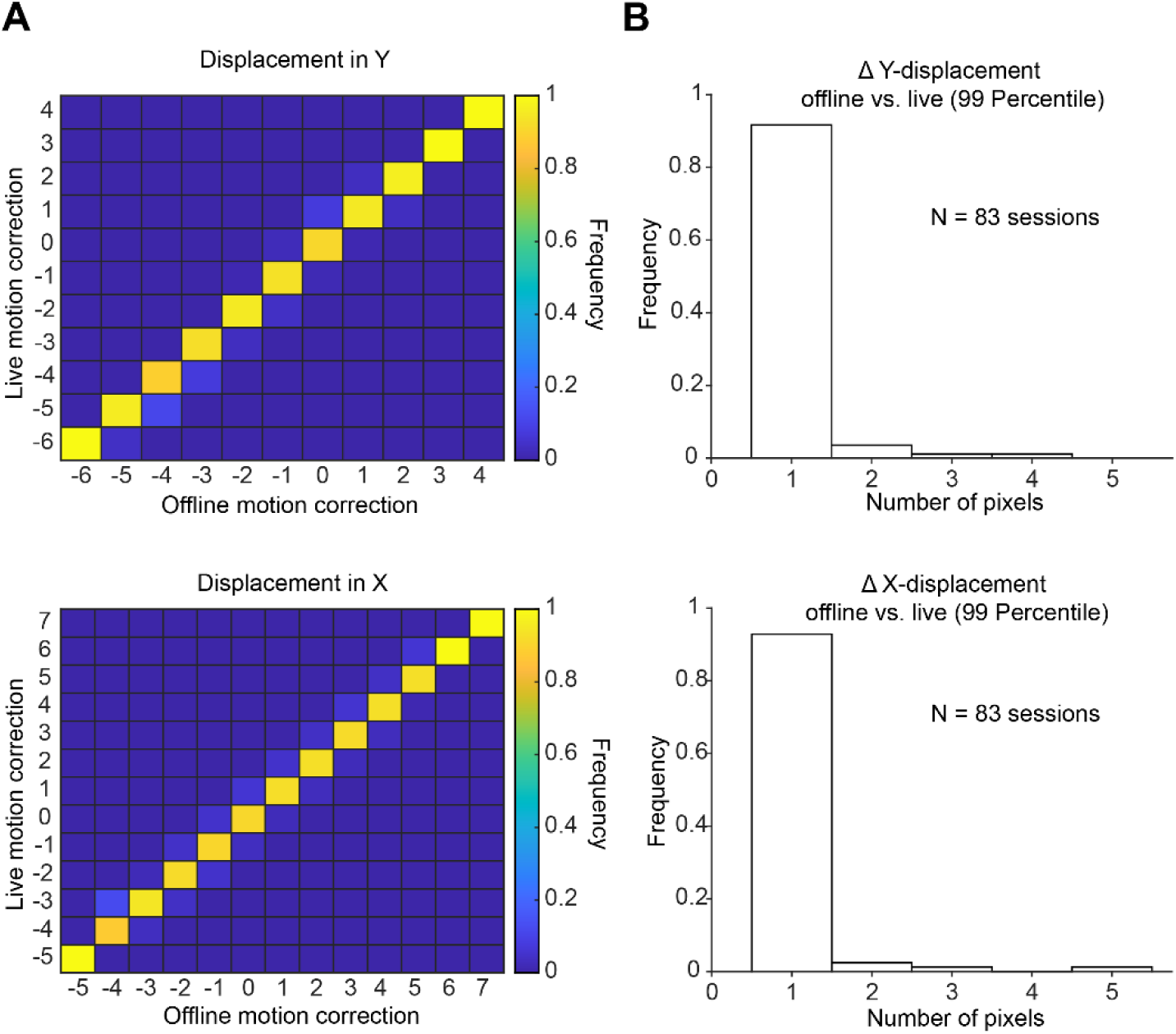
Validation of live motion correction a, Live motion correction performance for a given imaging session. We validated our live motion-correction algorithm by comparing its performance against a commonly used offline motion correction algorithm^65^. To be able to compare performance, we motion-corrected against the same reference image used for offline motion correction. For a given pixel displacement in X and Y in the offline algorithm, we calculated the proportion of frames which our online algorithm estimated to be displaced by the same amount. Each column is normalized to the total number of frames for a given displacement value. **b,** For X and Y displacements, in all our imaging sessions, the 99^th^ percentile of the difference between live and offline motion correction. Our data show that in more than 90% of our imaging sessions, the estimated pixel displacement using live and offline motion correction differ by 1 pixel at the 99^th^ percentile level.

**Supplementary Figure 3:**
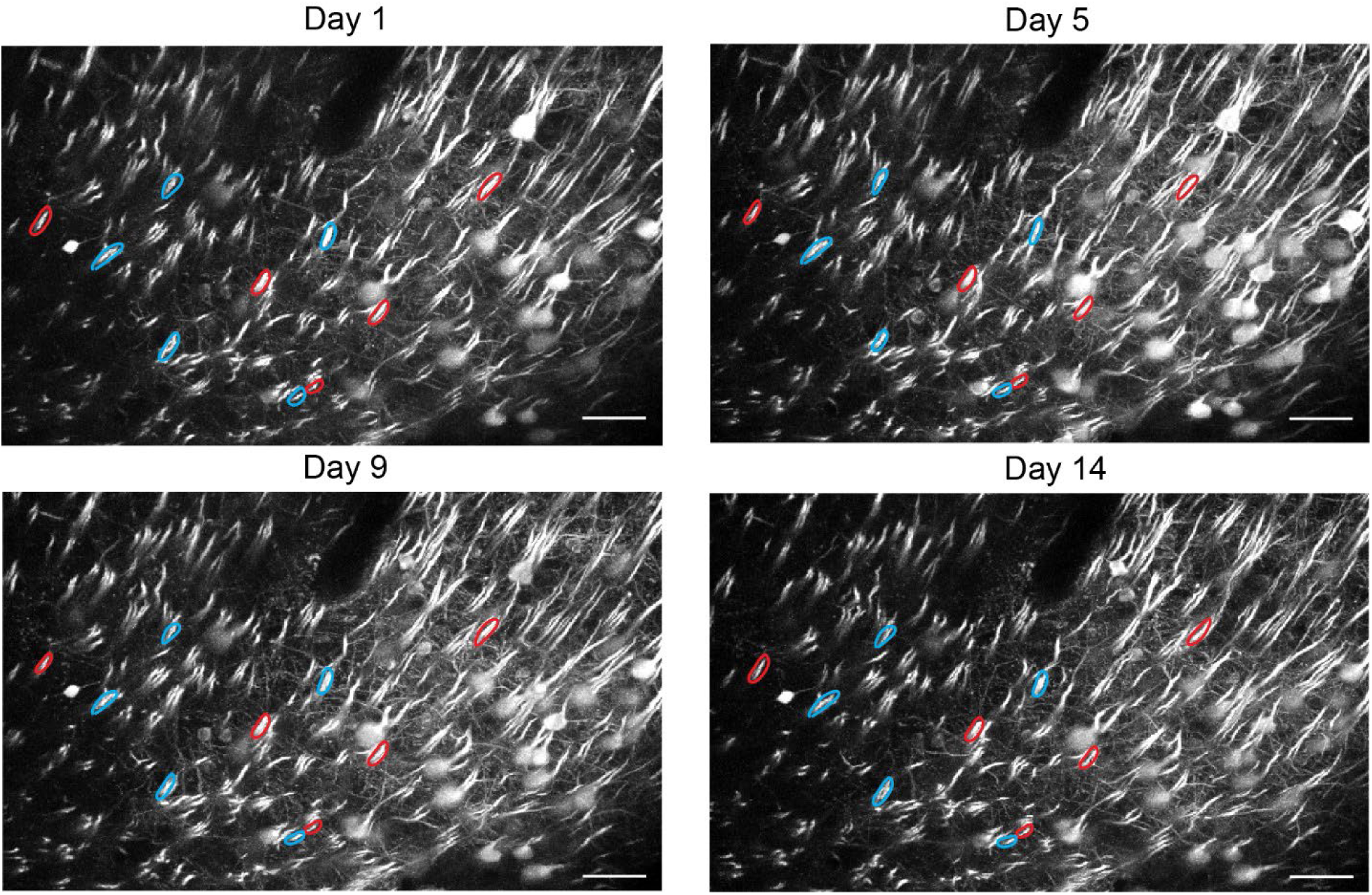
Representative field of view with P+ and P-neurons. A representative field of view with the same, GCaMP7f-labelled, chronically-tracked P+ and P- neurons (5 each, shown in red and blue, respectively), imaged at the proximal trunk (as a proxy for soma) during 14 days of learning, shown at day 1, 5, 9 and 14. P+ and P- neurons were selected on day 1. Scale bar represents 50 µm.

**Supplementary Figure 4:**
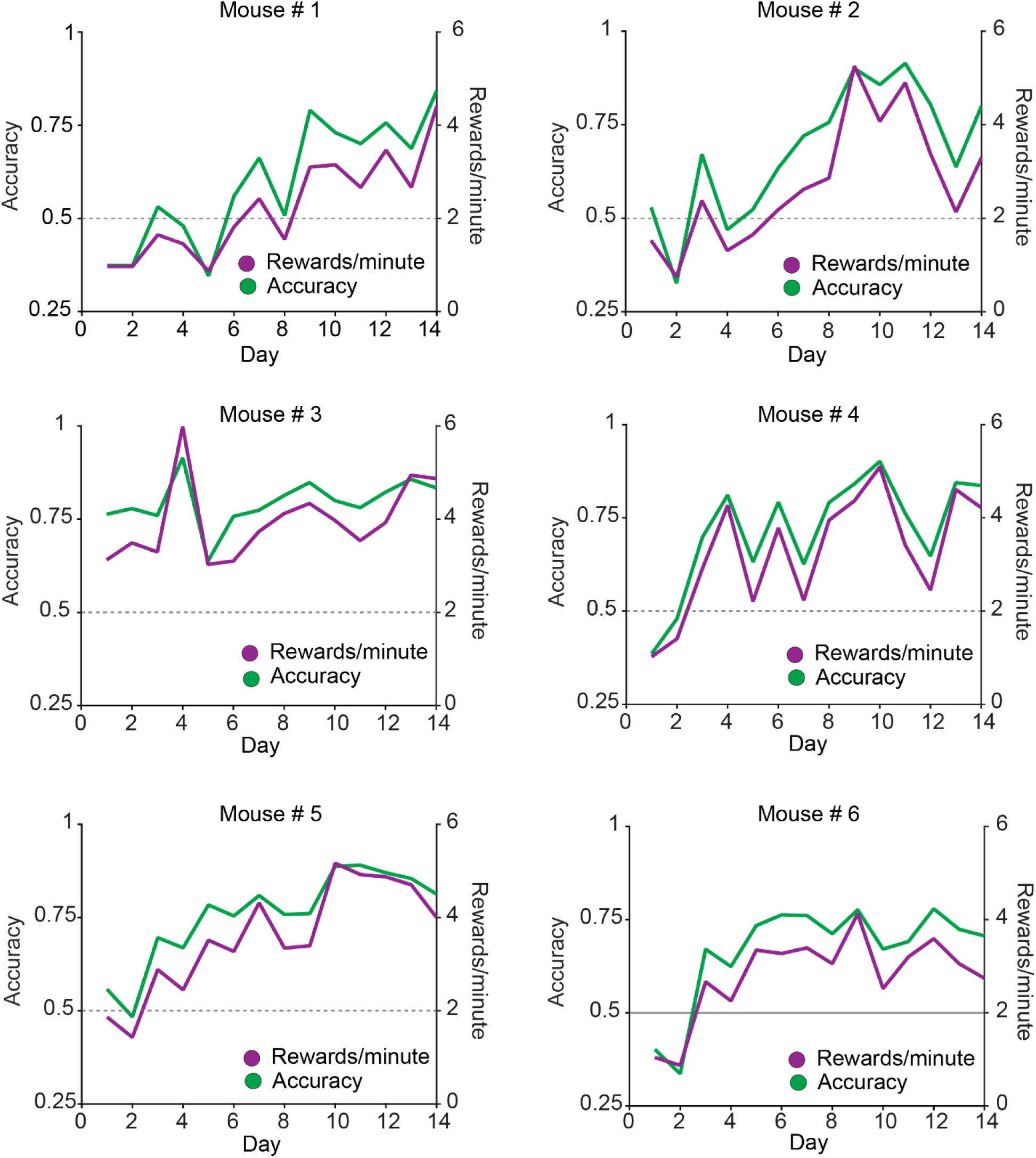
Task performance for individual animals. Task performance evaluated using two metrics: accuracy (the fraction of successful trials) and rewards per minute for each of 6 mice. Summary graph and statistics in Figure 1g. The grey dashed line represents chance accuracy.

**Supplementary Figure 5:**
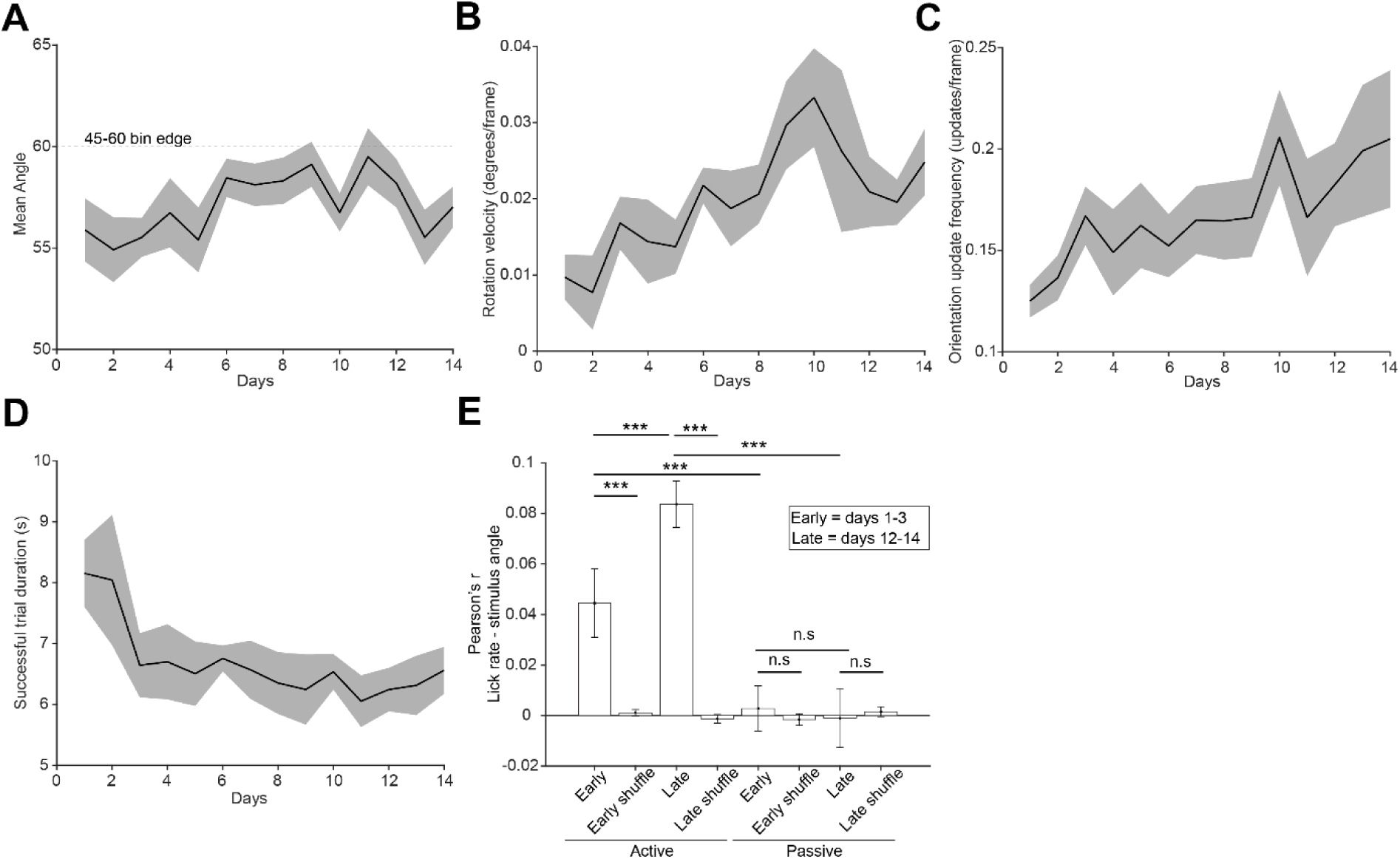
Visual stimulus and behavioral correlates of learning. a, Mean visual stimulus angle across 14 days of learning. Due to binning, stimuli below the 60-degrees threshold were presented as 45-degrees. **b,** Rotation speed across the 14 days of learning. Rotations towards and away from a reward were defined as positive and negative rotations, respectively. **c,** Mean frequency at which a new orientation was presented. **d,** Mean duration of successful trials across the 14 days of learning. **e,** Pearson’s correlation between licking frequency and stimulus angle (outside reward periods) during active (closed-loop) and passive stimulus presentation at the beginning and the end of learning (Two-way repeated measures ANOVA, p = 0.13, 1.4e^-11^ and 0.028 for the effect of state (active vs passive), days, and interaction between state and days, respectively. After two-stage Benjamini, Krieger and Yekutieli correction for false discovery rate, p = 8.5e^-4^ for early-active vs late-active, 3.4e^-4^ for early-active vs early-active (shuffle), 8.4e^-11^ for late-active vs late-active(shuffle), 1.7e^-4^ for early-active vs early passive, 5e^-11^ for late-active vs late-passive, 0.83 for early-passive vs late-passive, 0.8 for early-passive vs early-passive(shuffle), 0.81 for late-passive vs late-passive (shuffle)).

**Supplementary Figure 6:**
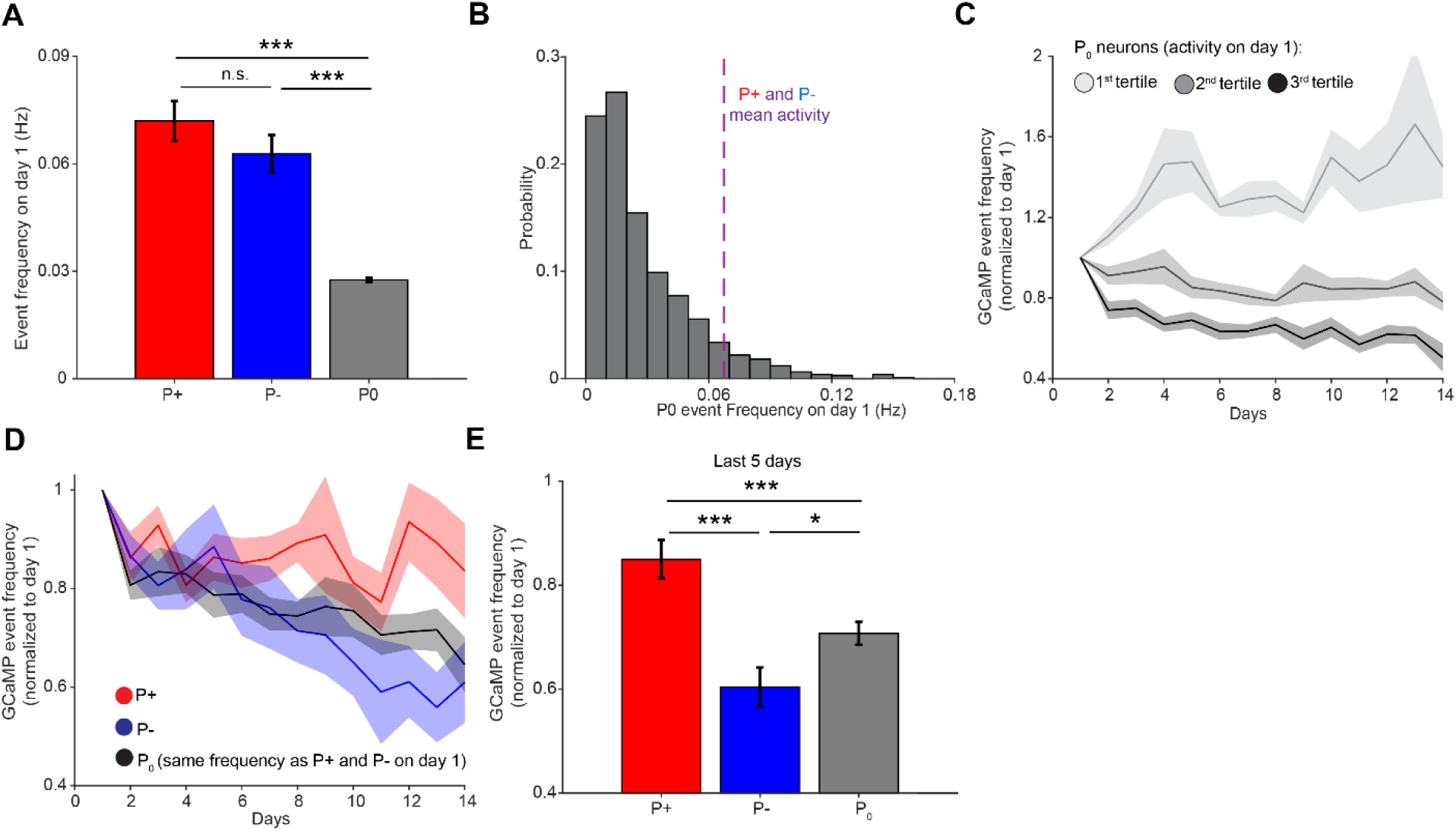
Changes in GCaMP transient frequency over days depend on day 1 frequency. a, Event frequency for P+, P- and P_0_ neurons on day 1. P+ and P- neurons were selected to be of higher activity on day 1 compared to other neurons in the field of view (One-way ANOVA, p = 2.52e^-30^. After Tukey’s multiple comparisons correction, p = 0.35, 9.57e^-10^ and 9.56e^-10^ for P+ vs P-, P+ vs P_0_ and P- vs P_0_, respectively. mean = 0.072, 0.063 and 0.028; SEM = 0.006, 0.005 and 5.6e^-4^; n = 27, 27 and 1839 for P+, P- and P_0_, respectively). **b**, Distribution of event frequency for P_0_ neurons on day 1. The mean event frequency for P+ and P- neurons (purple, dashed line) fall on the 8^th^ percentile of the P_0_ neurons distribution. **c,** Calcium transient frequency for P_0_ neurons divided in tertiles based on activity on day 1, across the 14 days of training normalized to the activity on day 1. All neurons were tracked over the full 14 days of imaging (n = 6 mice). **d,** Calcium transient frequency for P+, P- and P_0_ neurons. Only P_0_ neurons with the same calcium transient frequency as P+ and P- neurons on day 1 (neurons within the 95% confidence interval of the joint P+ and P- distribution) were selected. Neurons were tracked across the 14 days of imaging and activity was normalized to the activity on day 1 (Two-way repeated measures ANOVA, p = 0.008, p = 0.006 and p = 0.003 for the effect of population identity, days and an interaction between these 2, respectively. n = 6 mice across 14 days). **e,** Calcium transient frequency for P+, P- and P_0_ on days 10 to 14, normalized to day 1 (One-way repeated measures ANOVA, p = 1e^-7^. After Tukey’s multiple comparison, p = 3.3e^-6^, 5.47e^-4^ and 0.02 for P+ vs P-, P+ vs P_0_ and P- vs P_0,_ respectively; mean = 0.85, 0.60 and 0.71, SEM = 0.037, 0.037 and 0.02 for P+, P- and P_0_, respectively. n = 6 mice).

**Supplementary Figure 7:**
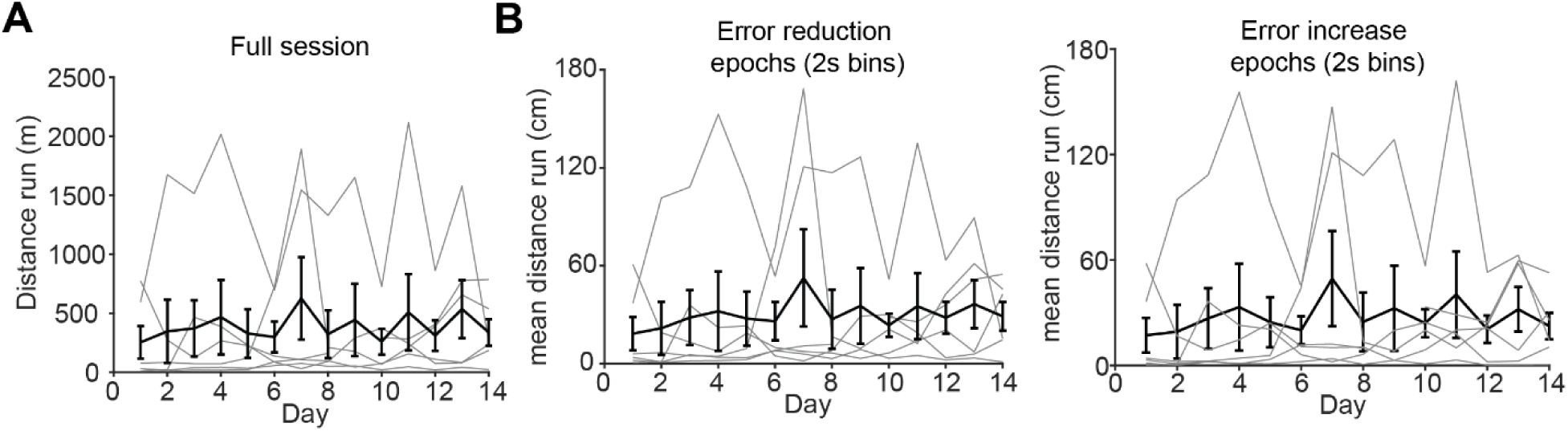
Learning is not due to changes in locomotion across days. a, In black, the mean and standard error of the mean for the distance run by all animals in individual imaging sessions. All sessions were of equal length (41.26 minutes or 76500 imaging frames (2 planes at 30Hz)). In grey, the distance run by individual mice (One-way repeated measures ANOVA, p = 0.58, n = 6 across 14 days). **b,** After excluding all timeouts, grey, black and reward-related periods, we divided our trials into 2 seconds bins and estimated whether the net rotation of the stimulus (the mean of the derivative angle) was positive (lefts panel, towards the reward, error reduction) or negative (right panel, away from the reward, error increase) in each bin and calculated the average distance run during error reduction and error increase epochs. We found no differences neither within condition across days, nor among conditions (Two-way repeated measures ANOVA, p = 0.15, 0.86 and 0.14 for the effects of error type, days and error type and days interaction n = 6 animals across 14 days).

**Supplementary Figure 8:**
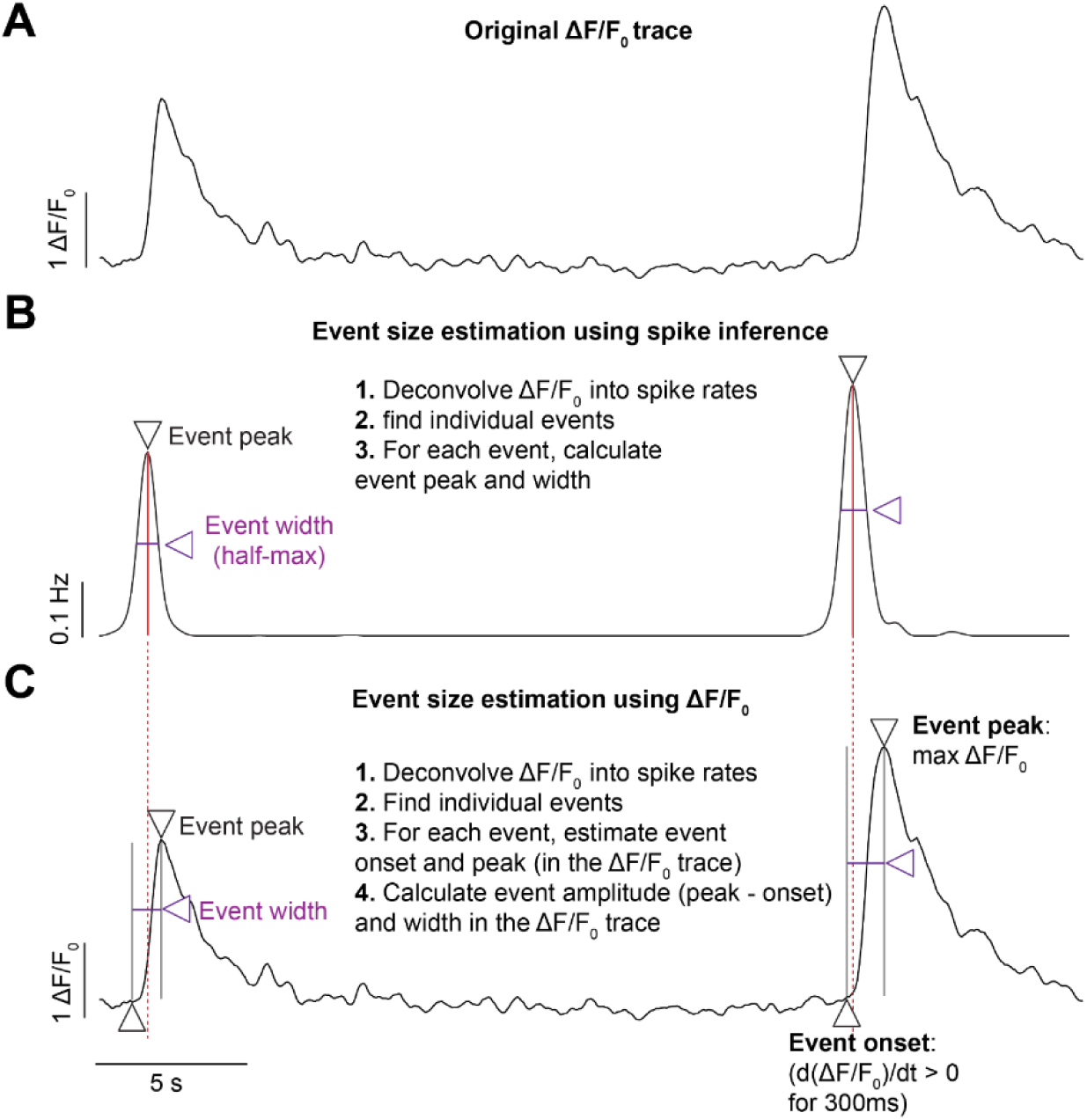
Event detection. a, ΔF/F_0_ trace for an individual neuron. **b,** For the same ΔF/F_0_ trace shown in A, calcium signal was deconvolved into a spike rate using CASCADE^44^. Event size was calculated by first detecting individual peaks (downward facing triangle, *findpeaks* function in MATLAB, see Method) and then calculating the area under the curve by estimating the amplitude (red, vertical line) and width at half maximum (purple line). **c,** Same ΔF/F_0_ trace as in a. To obtain ΔF/F_0_ metrics of event size, we first deconvolved ΔF/F_0_ signals into spike rates using CASCADE and detected individual transients (same as steps 1-2 as in b). For each detected event we then found its start (upward facing triangle) and peak (downward facing triangle) as the point at which the derivative of ΔF/F_0_ trace became positive for 300 ms in a row in a 2 seconds backward window from the detected spike inference peak, and the maximum ΔF/F_0_ value in the following 2 seconds (unless constrained by the presence of another event). The amplitude of the event was defined as ΔF/F_0_ value at peak – ΔF/F_0_ at onset. The width was defined as peak – onset frame.

**Supplementary Figure 9:**
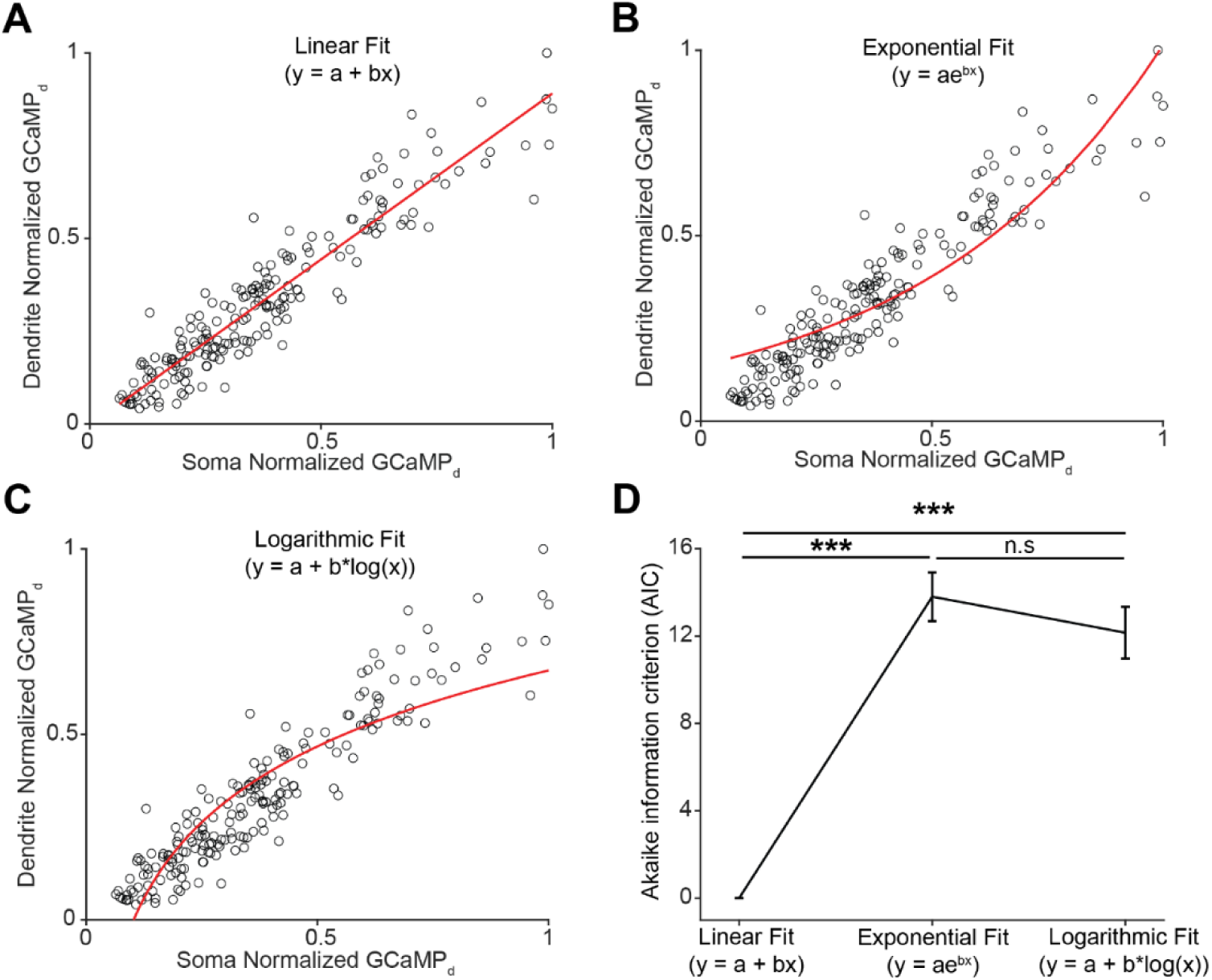
Testing the assumption of linearity between somatic and dendritic event magnitude. a, Representative example of a single neuron whose events were isolated and their magnitude estimated in the soma and dendrites, plotted against each other. Axes are normalized to the largest event. To describe the relationship between somatic and dendritic event magnitude, we fit a linear model. **b,** Same as in a, except this time we fit an exponential model to the data. **c,** Same as in a and b, this time however we fit a logarithmic model to the data. **d,** For all neurons, we calculated the Akaike information criteria (AIC) to describe the goodness of a model (while penalizing for the number of parameters used). Lower values of AIC mean better fit. AIC values were zeroed compared to the linear fit model. Error bars represent standard error of the mean (One-way repeated measures ANOVA; p < 1e^-15^, p < 1e^-15^, p = 0.56 for Linear vs Exponential, Linear vs Logarithmic and Exponential vs. Logarithmic after Tukey’s multiple comparison correction. Mean = 0, 13.8 and 12.1; SEM = 0, 1.11 and 1.17; n = 543 neurons for Linear, Exponential and logarithmic fit, respectively).

**Supplementary Figure 10:**
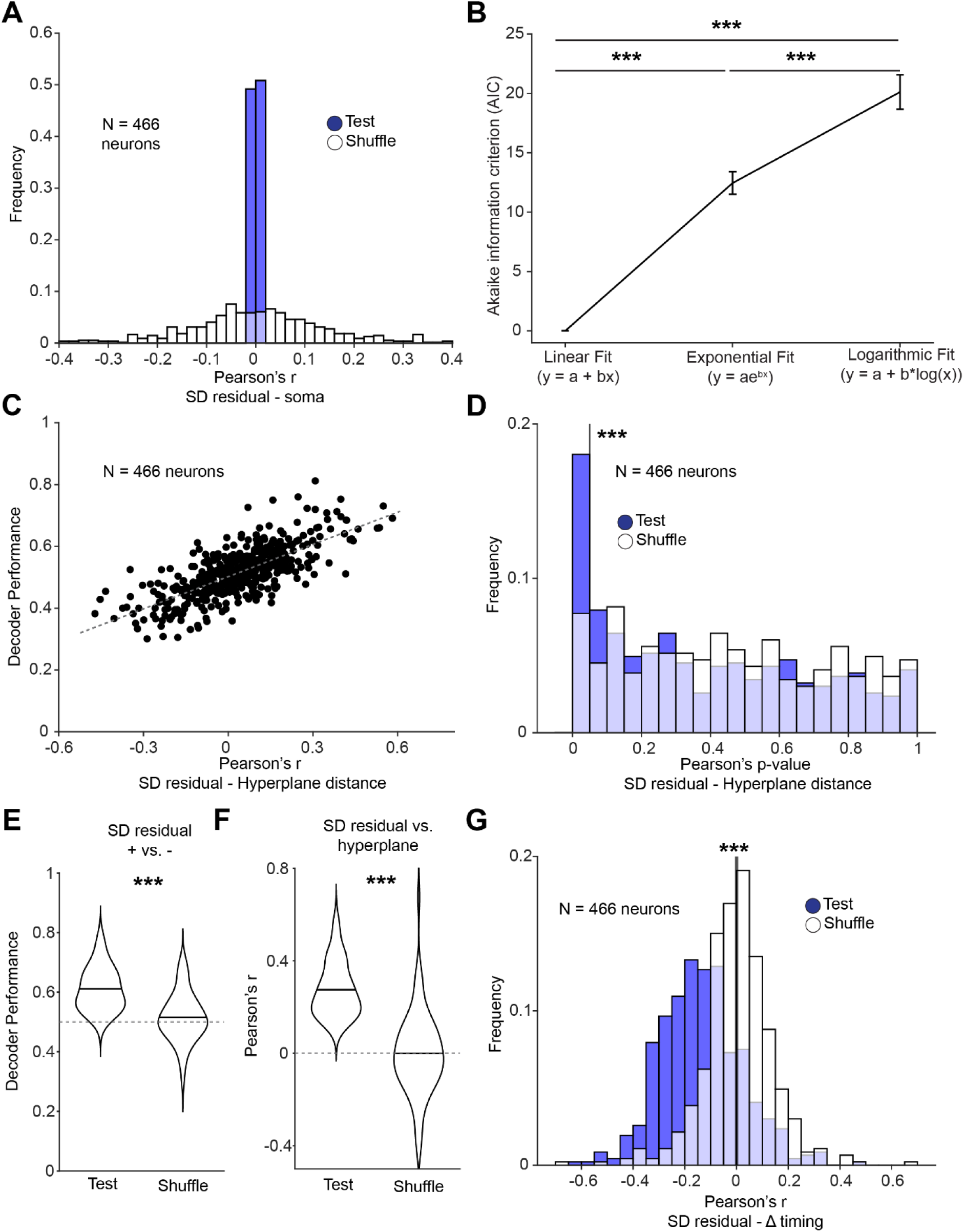
Decoding somato-dendritic residuals using ΔF/F_0_-based event size estimation. a, Same as in Figure 2i. Our ΔF/F_0_-based approach to estimate event size (see Methods and Supplementary Figure 2) decorrelated the SD residual from somatic magnitude. **b,** Same as Supplementary Figure 1. Linear models best describe the relationship between somatic and dendritic event magnitudes. For all neurons, we calculated the Akaike information criteria (AIC). Lower values of AIC mean a better fit. AIC values were zeroed compared to the linear fit model. Error bars represent standard error of the mean (One-way repeated measures ANOVA; p = 3e^- 10^, p = 1e^-10^, p = 1e^-10^ for Linear vs Exponential, Linear vs Logarithmic and Exponential vs. Logarithmic after Tukey’s multiple comparison correction. Mean = 0, 12.4 and 20.1; SEM = 0, 0.9 and 1.4; n = 543 neurons for Linear, Exponential and logarithmic fit, respectively). **c,** Same as Figure 2g. Pearson correlation between decoder performance (linear SVM) and the correlation between SD residual and the distance from the hyperplane (or SVM classification confidence). Each dot represents one neuron (Pearson correlation = 0.73. p = 2.5e^-78^). **d,** Same as Figure 2h, the proportion of neurons with a statistically-significant (alpha = 0.05) correlation between the SD residual and the distance from the hyperplane (or classification confidence, Wilcoxon signed rank test = p = 7.3e^-4^. N = 466 neurons). **e,** Same as Figure 2j. Decoder performance (linear SVM) for statistically-significant neurons in d (paired t-test, p = 8.3e^-10^. Mean = 0.61 and 0.51; SEM = 0.008 and 0.01; n = 66). **f,** Same as Figure 2k. Pearson correlation between the SD residual and the distance from hyperplane for statistically-significant neurons (paired t-test, p = 1.6e^-15^. Mean = 0.27 and-9.6e^-4^; SEM = 0.01 and 0.02; n = 66). **g,** Same as Figure 2m. Pearson correlation between relative event timing in soma and dendrites and SD residual (residual from best fit). A negative correlation means that the larger the residual (more dendritically-amplified) the earlier the peak timing in the dendrite compared to the soma, paired t-test, p = 9.7e^-40^. Mean = 0.13 and 0.005; SEM = 0.007 and 0.006; n = 466).

**Supplementary Figure 11:**
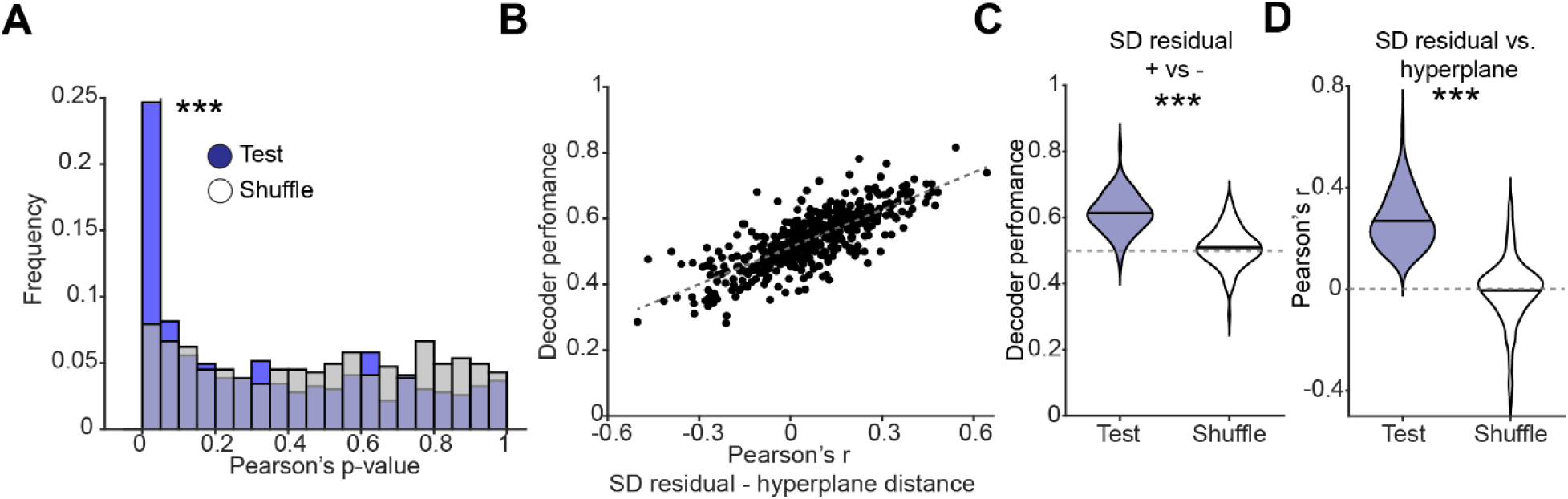
Decoding SD residuals using network activity in a single frame preceding event onset. a, Decoder performance as a function of the correlation between SD residuals and hyperplane distance (Pearson’s r = 0.77; p-value = 7.1e^-94^, n = 466 neurons). Datapoints represents individual neurons. **b,** Distribution of p-values for test data and a control randomly shuffled distribution, testing the correlation between SD residuals and distance from the hyperplane (or classification confidence, Wilcoxon signed rank test = p = 2.8e^-9^. N = 466 neurons). **c,** Decoding performance for neurons with a statistically significant correlation between SD residual and distance from the hyperplane (paired t-test, p = 8.6e^-28^. Mean = 0.61 and 0.49; SEM = 0.006 and 0.007; n = 99). Dashed grey line indicates chance level. **d,** Pearson’s r for neurons with a statistically-significant correlation between SD residual and the distance between population vector and hyperplane (paired t-test, p = 7.9e^-30^. Mean = 0.27 and-0.005; SEM = 0.01 and 0.01; n = 99).

**Supplementary Figure 12:**
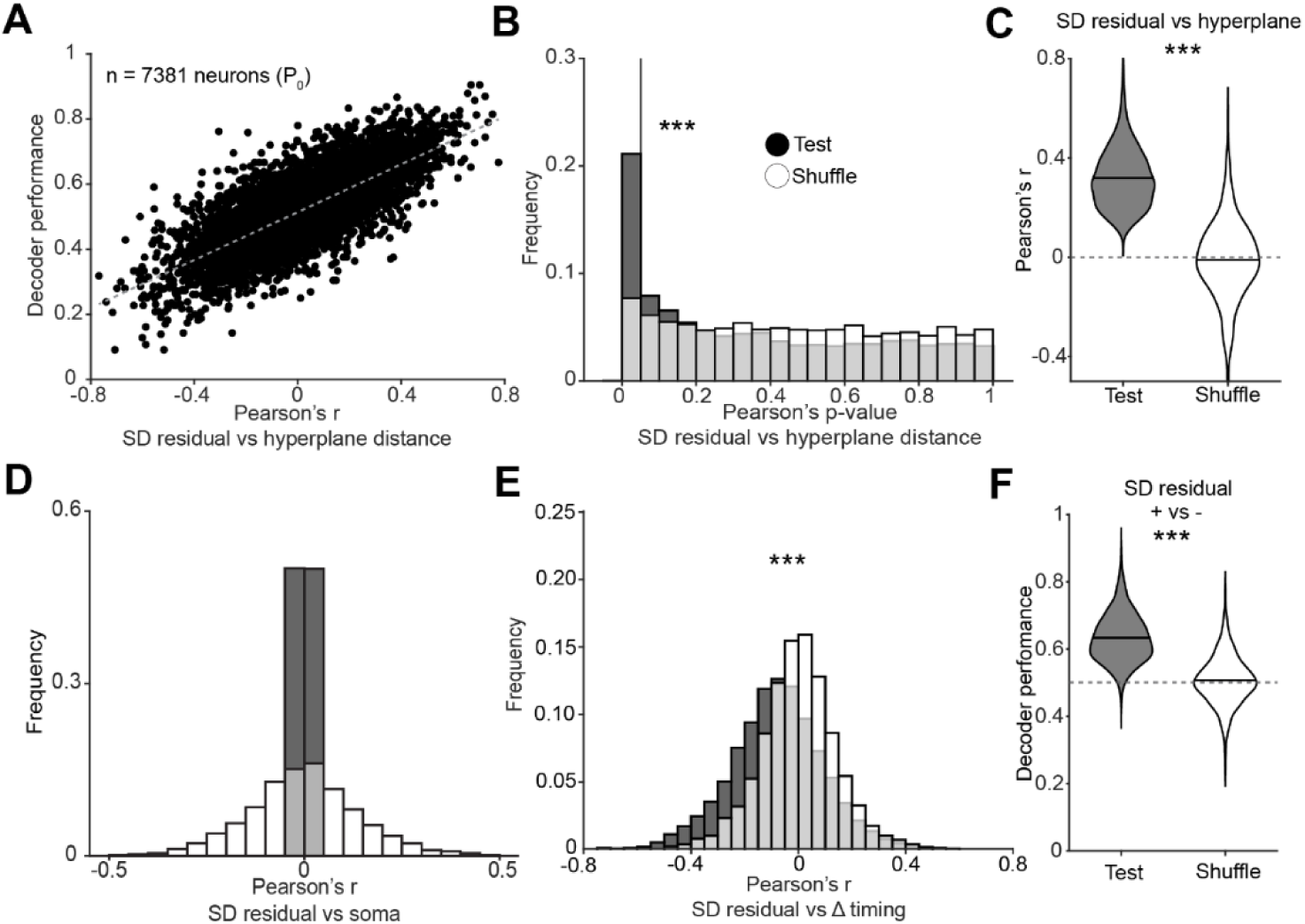
Differences in somatic and dendritic magnitudes for coincident events are predicted by local network dynamics in P_0_. a, Decoder performance as a function of the correlation between SD residuals and hyperplane distance (Pearson’s r = 0.75; p-value < 2.2e^-308^, n = 7381 neurons). Datapoints represents individual neurons. **b,** Distribution of p-values for test data and a control randomly shuffled distribution, testing the correlation between SD residuals and distance from the hyperplane (or classification confidence, Wilcoxon signed rank test = p = 5e^-96^. N = 7381 neurons). **c,** Pearson’s r for neurons with a statistically-significant correlation between SD residual and the distance between population vector and hyperplane (paired t-test, < 2.2e^-308^. Mean = 0.31 and-0.01; SEM = 0.003 and 0.005). **d,** For all neurons, the Pearson’s r for somato-dendritic residuals and somatic event magnitude. **e,** Pearson correlation value between the SD residual the event latency between soma and its corresponding dendrite indicating that the larger the SD residual, the earlier the dendritic peak is compared to the somatic one (paired t-test, p = 1.48e^-169^. Mean =-0.077 and-6.9e^-4^; SEM = 0.002 and 0.001; n = 7381 neurons). **f,** Decoding performance for neurons with a statistically significant correlation between SD residual and distance from the hyperplane (paired t-test, p = 1.1e^-254^. Mean = 0.63 and 0.50; SEM = 0.002 and 0.002). Dashed grey line indicates chance level.

**Supplementary Figure 13:**
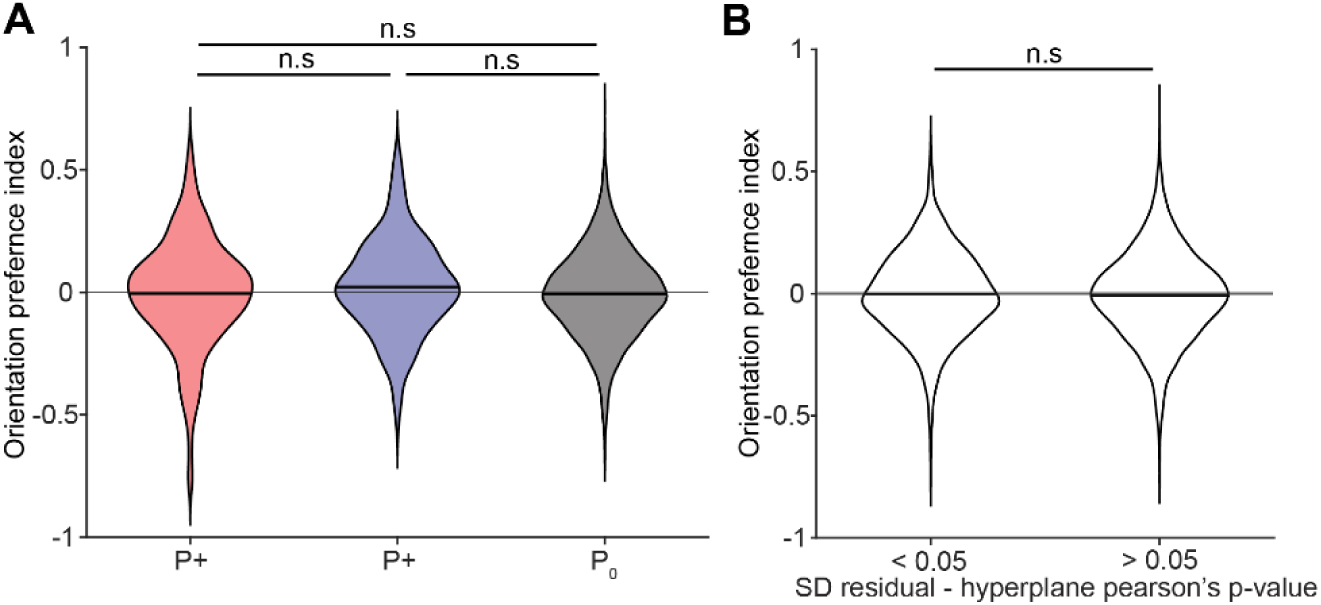
Orientation preferences of decoded neurons. a, Orientation preference index, defined as (R_90_ – R_0_)/ (R_90_ + R_0_) where R_90_ and R_0_ are the inferred spike rates at 90-and 0-degrees angles, respectively during passive visual stimulus presentation for P+, P- and P_0_ (One-way ANOVA, p = 0.09; After Tukey’s multiple comparisons p = 0.25, 0.98 and 0.07 for P+ vs P-, P+ vs P_0_ and P- vs P_0_, respectively. Mean =-0.005, 0.02 and-0.007; SEM = 0.01, 0.01 and 002; n = 268, 198 and 7381 for P+, P- and P_0_, respectively). **b,** Orientation preference index for all neurons with a statistically-significant Pearson’s correlation between the SD residual and the network distance from the maximally-separating hyperplane (t-test, p = 0.31. Mean =-0.002 and-0.007; SEM = 0.004 and 0.002; n = 1298 and 6547 for significant and non-significant neurons, respectively).

**Supplementary Figure 14:**
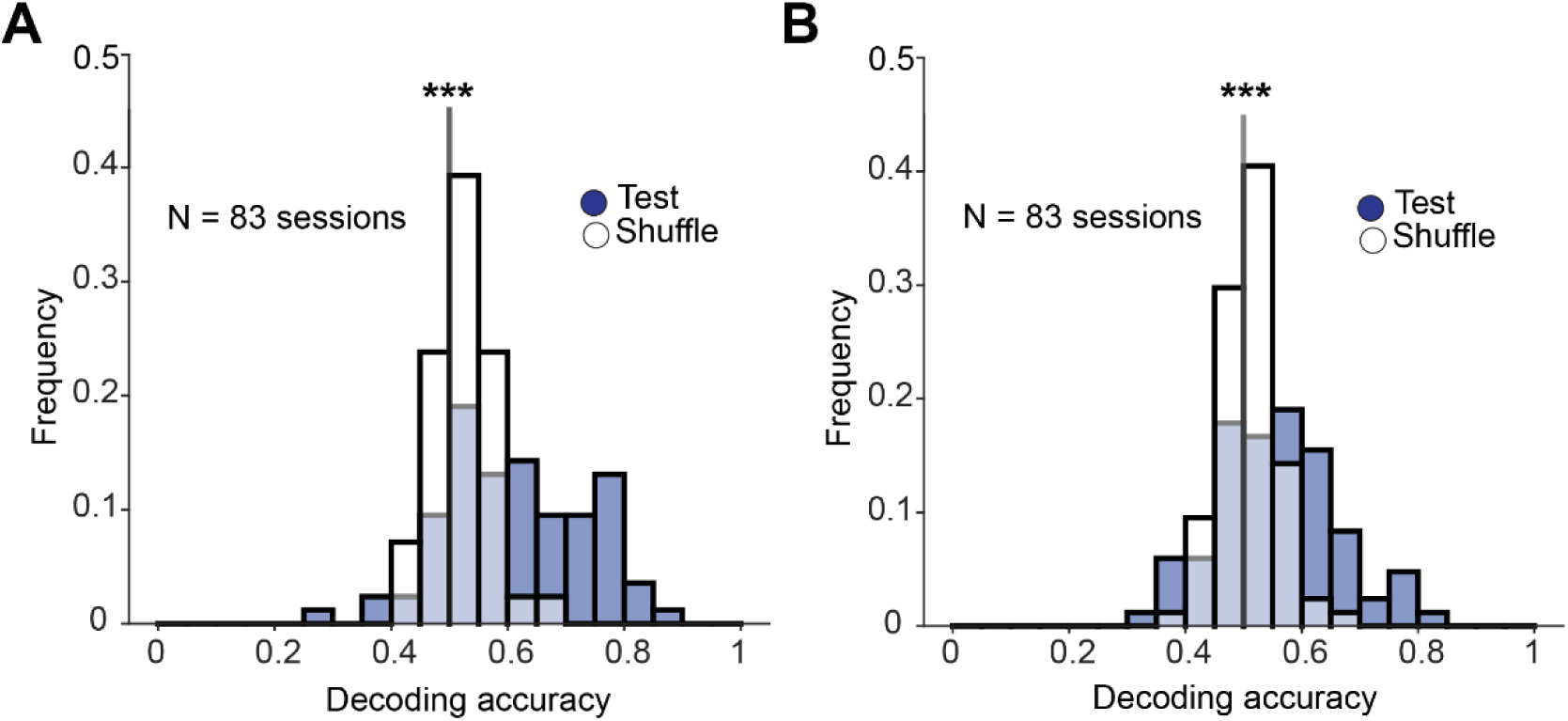
Decoding task related variables using ΔF/F_0_-based event size estimation. a, b, Same as Figure 3e (left panel, paired t-test, p = 4.3e^-9^. Mean = 0.61 and 0.52; SEM = 0.01 and 0.01, respectively for test and shuffle data. N = 83 sessions) and 3g (right panel, paired t-test, p = 2.4e^-4^. Mean = 0.55 and 0.50; SEM = 0.01 and 0.01, respectively for test and shuffle data. N = 83 sessions) respectively, using a ΔF/F_0_-based metrics for estimating the size of the SD residual.

**Supplementary Figure 15:**
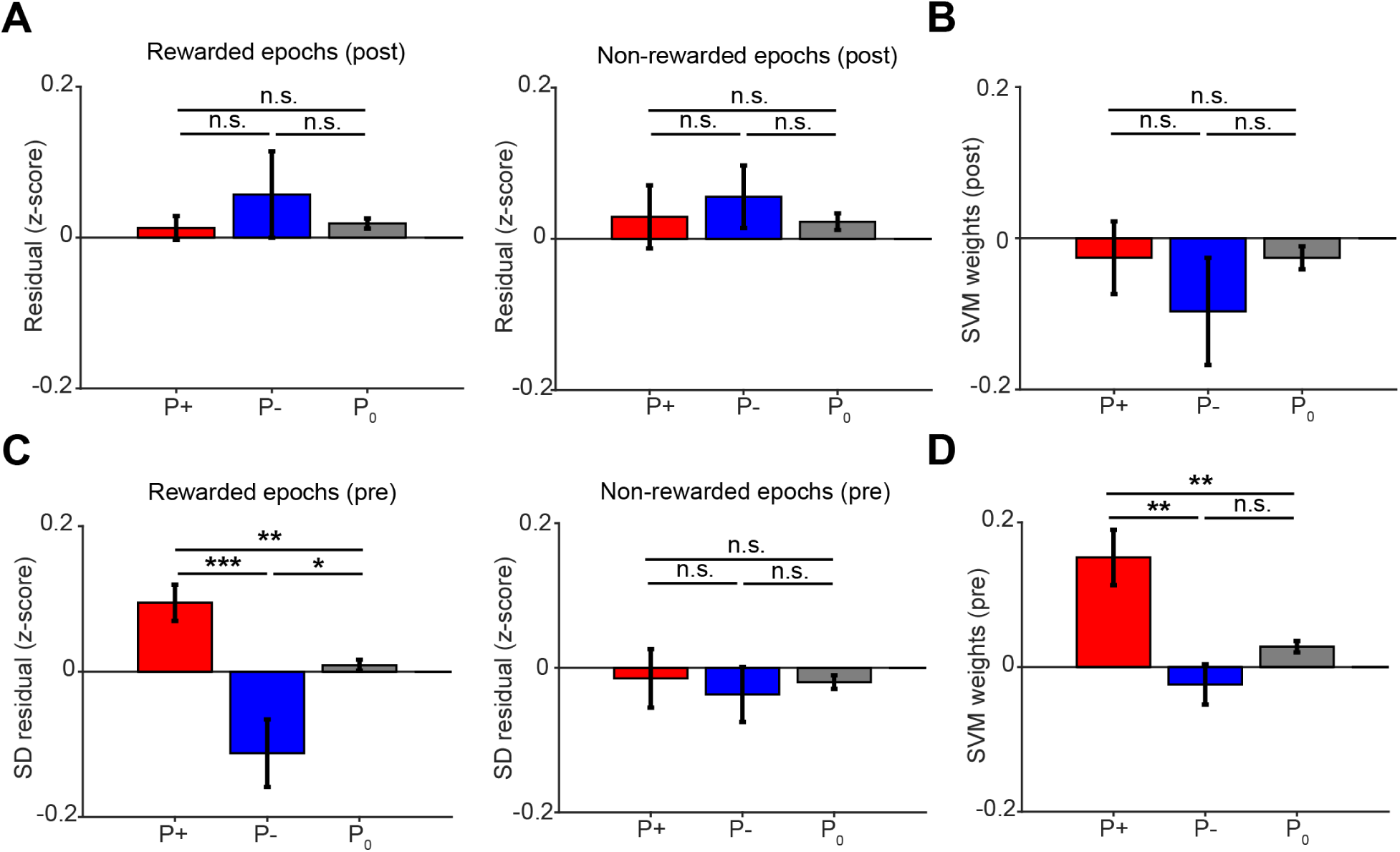
Dendritic reward signals are cell-specific during periods of reward anticipation but not during reward consumption. a, SD residuals in the two seconds following target reach for rewarded trials (left) and the end of a trial for rewarded and unrewarded trials (right), respectively, for P+, P- and P_0_ neurons (rewarded epochs, One-way repeated measures ANOVA, p = 0.5. After Tukey’s multiple comparison p = 0.75, p = 0.93 and p = 78 for P+ vs P-, P+ vs P_0_ and P- vs P_0_, respectively. Mean = 0.01, 0.06 and 0.02; SEM = 0.02, 0.06 and 0.01 for P+, P- and P_0,_ respectively. Unrewarded epochs, One-way repeated measures ANOVA, p = 0.47. After Tukey’s multiple comparison p = 0.65, p = 0.99 and p = 0.57 for P+ vs P-, P+ vs P_0_ and P- vs P_0_, respectively. Mean = 0.03, 0.06 and 0.02, SEM = 0.04, 0.04 and 0.01 for P+, P- and P_0,_ respectively. n = 84 sessions). **b,** SVM weights for decoding rewarded vs unrewarded trials after the end of a trial for P+, P- and P_0_ neurons. Here, the rewarded side of the hyperplane was arbitrarily assigned as the positive side, while the unrewarded one was assigned as the negative one (One-way repeated measures ANOVA, p = 0.72. After Tukey’s multiple comparison p = 0.87, p = 0.99 and p = 0.71 for P+ vs P-, P+ vs P_0_ and P- vs P_0_, respectively. Mean =-0.03,-0.1 and-0.03; SEM = 0.05, 0.07 and 0.2 for P+, P- and P_0,_ respectively. n = 84 sessions). **c,** SD residual in the two seconds preceding target reach for rewarded trials (left) and in the two seconds preceding the end of a trial for unrewarded trials (right), for P+, P- and P_0_ neurons (rewarded epochs, One-way repeated measures ANOVA, p = 2e^-4^. After Tukey’s multiple comparison p = 5.6e^-4^, p = 4.8e^-3^ and p = 0.035 for P+ vs P-, P+ vs P_0_ and P- vs P_0_, respectively. Mean = 0.09,-0.11 and 0.01, SEM = 0.03, 0.05 and 0.01 for P+, P- and P_0,_ respectively. Unrewarded epochs, One-way repeated measures ANOVA, p = 0.83. After Tukey’s multiple comparison p = 0.92, p = 0.99 and p = 0.9 for P+ vs P-, P+ vs P_0_ and P- vs P_0_, respectively. Mean =-0.01,-0.04 and-0.01, SEM = 0.04, 0.04 and 0.01 for P+, P- and P_0,_ respectively. n = 84 sessions). **d,** SVM weights for decoding rewarded vs unrewarded trials for P+, P- and P_0_ neurons. As in b, the rewarded side of the hyperplane was arbitrarily assigned as the positive side, while the unrewarded one was assigned as the negative one (One-way repeated measures ANOVA, p = 4.1e^-4^. After Tukey’s multiple comparison p = 3.8e^-3^, p = 5.6e^-3^ and p = 0.18 for P+ vs P-, P+ vs P_0_ and P- vs P_0_, respectively. Mean = 0.15,-0.02 and 0.03, SEM = 0.04, 0.03 and 0.01 for P+, P- and P_0,_ respectively. n = 84 sessions).

**Supplementary Figure 16:**
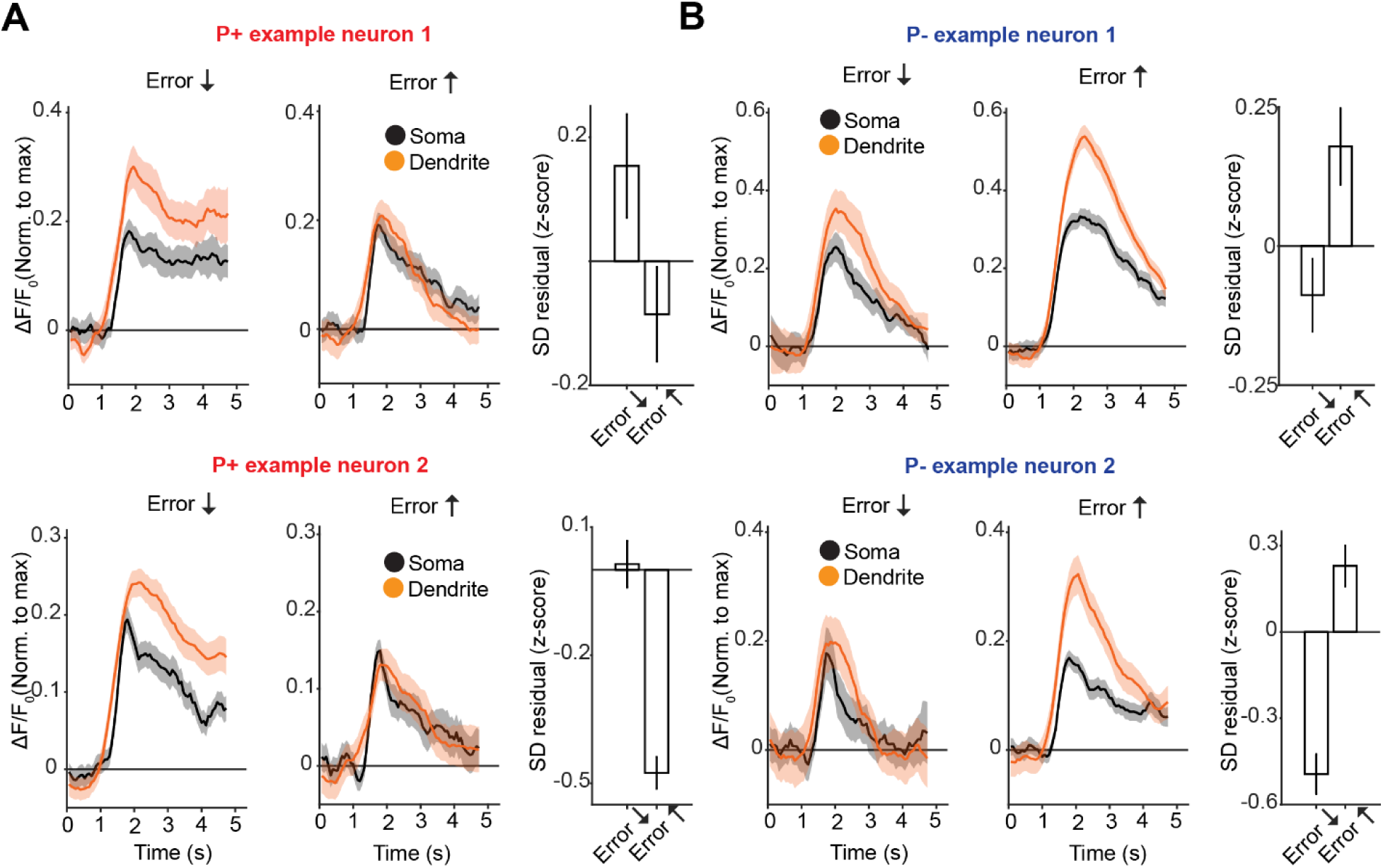
Four additional example neurons (2 P+ and 2 P-) showing different somato-dendritic relationships during error decrease and error increase epochs. For two P+ and two P- neurons, mean ΔF/F0 signal in the soma (black) and in the dendrite (orange) are shown for all events that occurred during epochs of error reduction and error increase, respectively. The bar graph represents the mean SD residual value (z-scored) for all events occurred during error decrease and increase epochs. Shaded areas and error bars represent SEM.

**Supplementary Figure 17:**
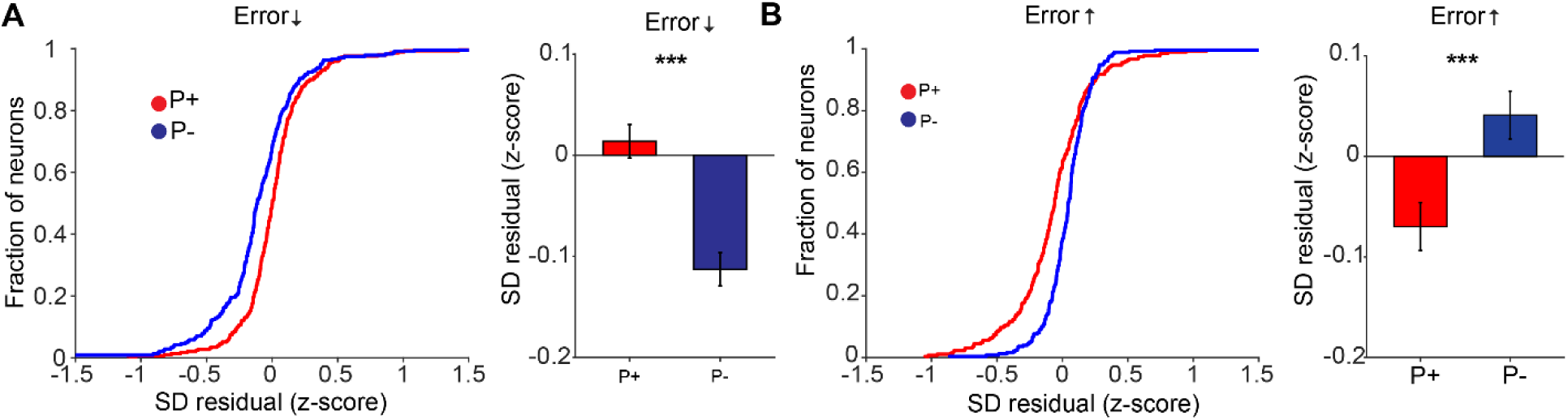
Neuron specific error signals using our ΔF/F_0_ metric to estimate event size. a, Same as Figure 4c. During error reduction epochs, dendrites of P+ neurons are relatively amplified compared to the dendrites of P- neurons (t-test, p = 1.4e^-5^; Mean = 0.01 and-0.11; SEM = 0.02 and 0.02; n = 292 and 240 for P+ and P- neurons, respectively). **b,** Same as Figure 4d. During error reduction epochs, dendrites of P+ neurons are relatively attenuated compared to the dendrites of P- neurons (t-test, p = 3.1e^-6^; Mean =-0.07 and 0.04; SEM = 0.02 and 0.01; n = 267 and 249 for P+ and P- neurons, respectively).

**Supplementary Figure 18:**
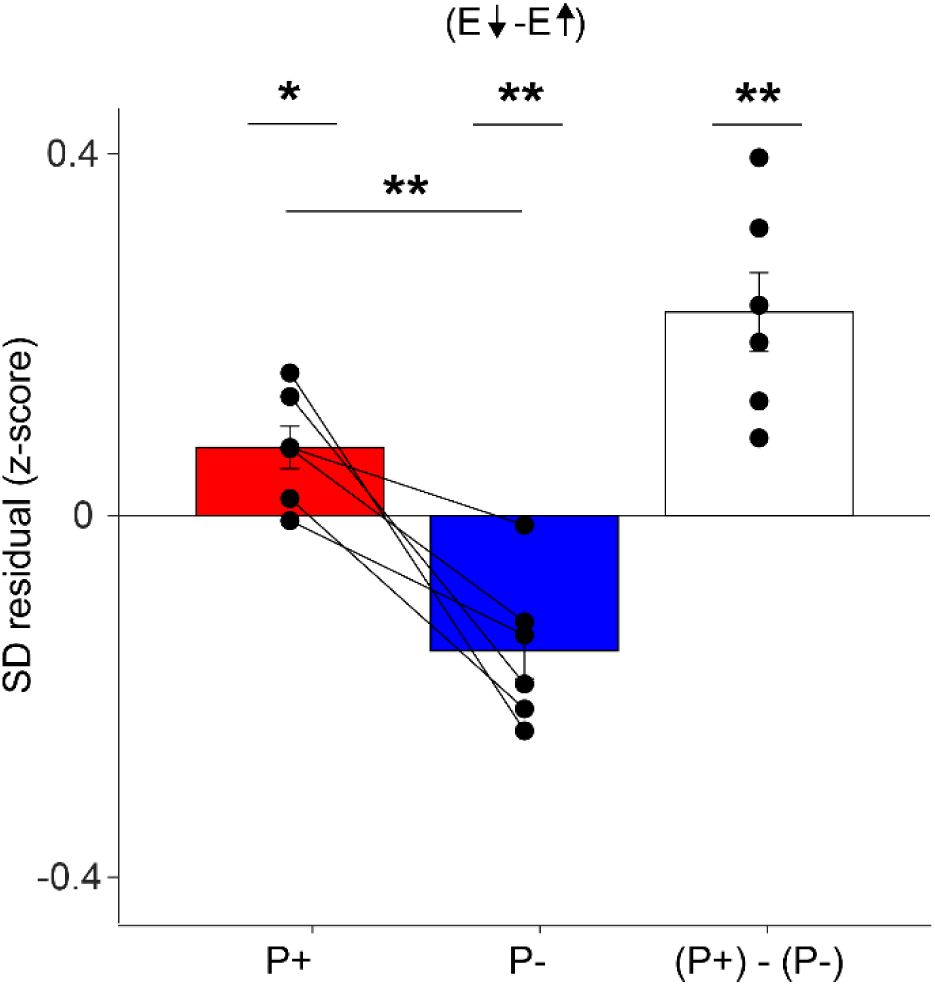
Dendritic signals are consistent across all mice. For all mice (N = 6) the separability of dendritic signals, defined as the SD residual during epochs of error decrease minus the SD residual during epochs of error increase, for P+ (in red), P- (in blue), and overall dendritic separability ((P+) – (P-)) (Paired t-test, p = 0.03, p = 0.006 and p = 0.005; Mean = 0.08,-0.15 and 0.23; SEM = 0.03, 0.03 and 0.05 for P+, P- and (P+) – (P-), respectively. N = 6 animals. Note that comparing P+ vs. P- or (P+) – (P-) vs. 0 is statistically equivalent).

**Supplementary Figure 19:**
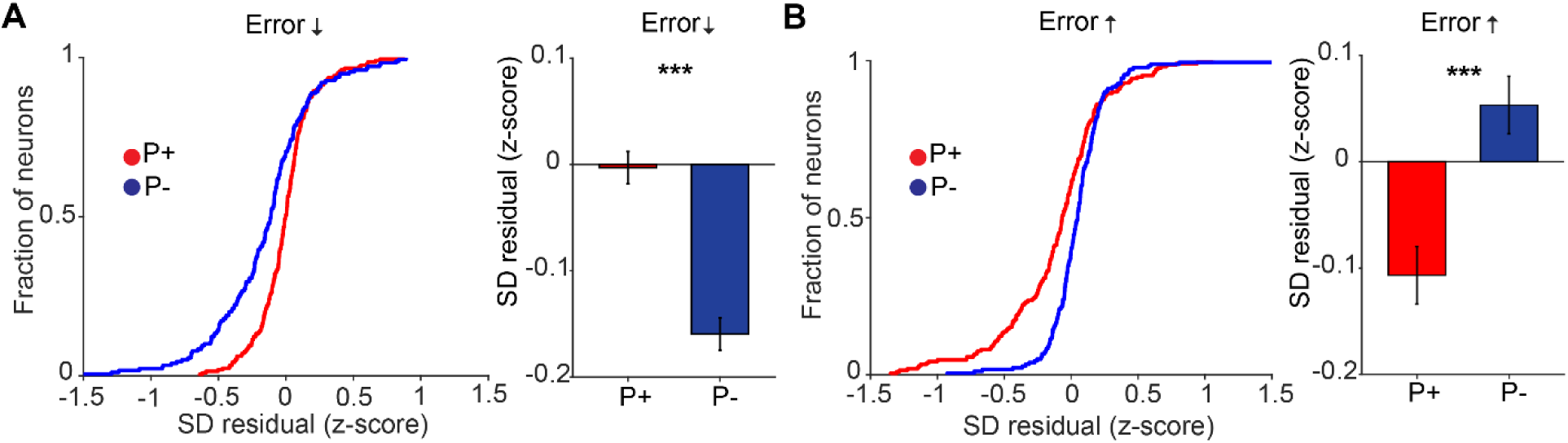
Dendritic error signals cannot be explained away by differences in somatic event magnitude. a, b, The cumulative distribution function for SD residuals (z-scored per neuron) in P+ (red) and P- (blue) neurons during error decrease in a (t-test, p = 3.5e^-6^, mean =-0.003 and-0.16; SEM = 0.015 and 0.03; n = 211 and 179 for P+ and P-) and error increase in b (t-test, p = 4.6e^-6^, mean - 0.1 and 0.05; SEM = 0.03 and 0.02; n = 211 and 179 for P+ and P- neurons) epochs. Only neurons with the same somatic response to error decrease and increase were selected for this analysis.

**Supplementary Figure 20:**
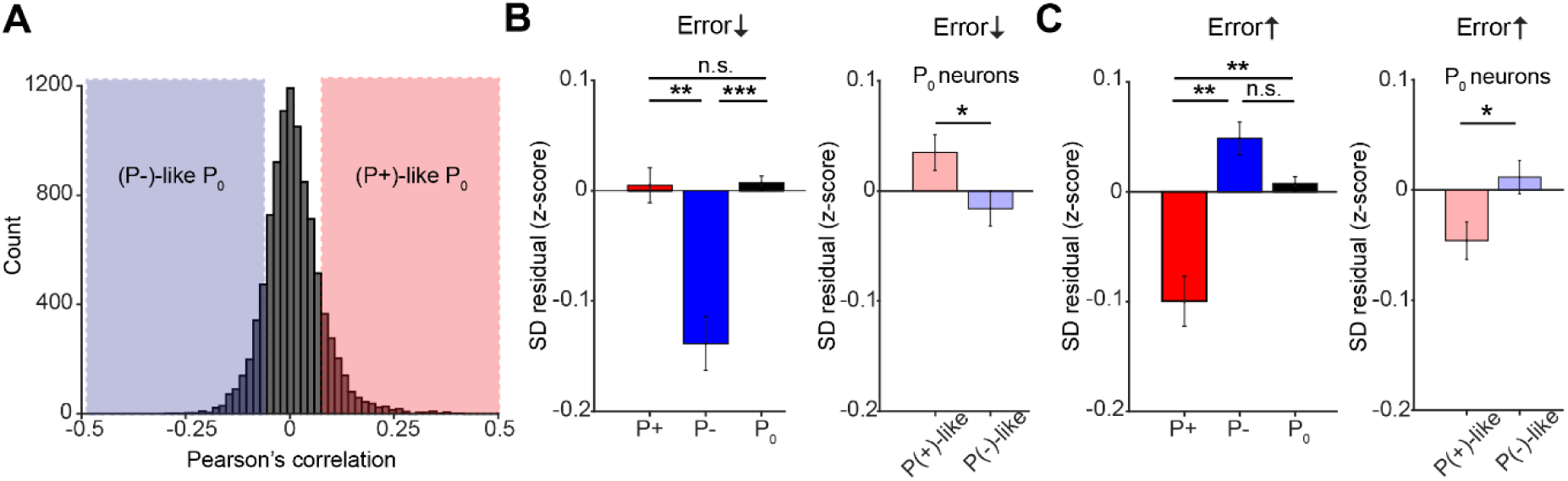
P_0_ neurons which are functionally-correlated to P+ and P- neurons also receive vectorized dendritic error signals. a, Across all sessions, we divided P_0_ neurons into (P+)- and P(-)-like based on their correlation to the (P+)-(P-) subtracted signal we used to rotate the visual stimulus. Among all P_0_ neurons (n = 9796) we defined (P+)-and (P-)-like P_0_ neurons as those with a correlation above 1 and below - 1 standard deviation. **b,** Left panel, during error reduction epochs, bar graph comparing the SD residual (z-scored) in P+, P-and P_0_ neurons (One-way ANOVA, p = 1.6e^-4^. After Tukey’s multiple comparisons, p = 0.006, p = 1 and p = 8.9e^-5^ for P+ vs P-, P+ vs P_0_ and P-vs P_0_, respectively. Mean = 0.004,-0.13 and 0.007; SEM = 0.02, 0.02 and 0.01; n = 292, 240 and 8335 for P+, P-and P_0_, respectively). Right panel, during error reduction epochs, bar graph comparing the SD residual (z-scored) in (P+)-and P(-)-like P_0_ neurons (t-test, p = 0.025; mean = 0.04 and-0.02; SEM = 0.02 and 0.02; n = 1059 and 1110 for (P+)-and P(-)-like P_0_ neurons, respectively). **c,** Left panel, during error increase epochs, bar graph comparing the SD residual (z-scored) in P+, P-and P_0_ neurons (One-way ANOVA, p = 0.002. After Tukey’s multiple comparisons, p = 0.004, p = 0.45 and p = 0.003 for P+ vs P-, P+ vs P_0_ and P-vs P_0_, respectively. Mean =-0.1, 0.05 and 0.01; SEM = 0.02, 0.02 and 0.01; n = 267, 249 and 8356 for P+, P-and P_0_, respectively). Right panel, during error increase epochs, bar graph comparing the SD residual (z-scored) in (P+)-and P(-)-like P_0_ neurons (t-test, p = 0.010; mean =-0.05 and 0.01; SEM = 0.02 and 0.01; n = 1070 and 1099 for (P+)-and P(-)-like P_0_ neurons, respectively).

**Supplementary Figure 21:**
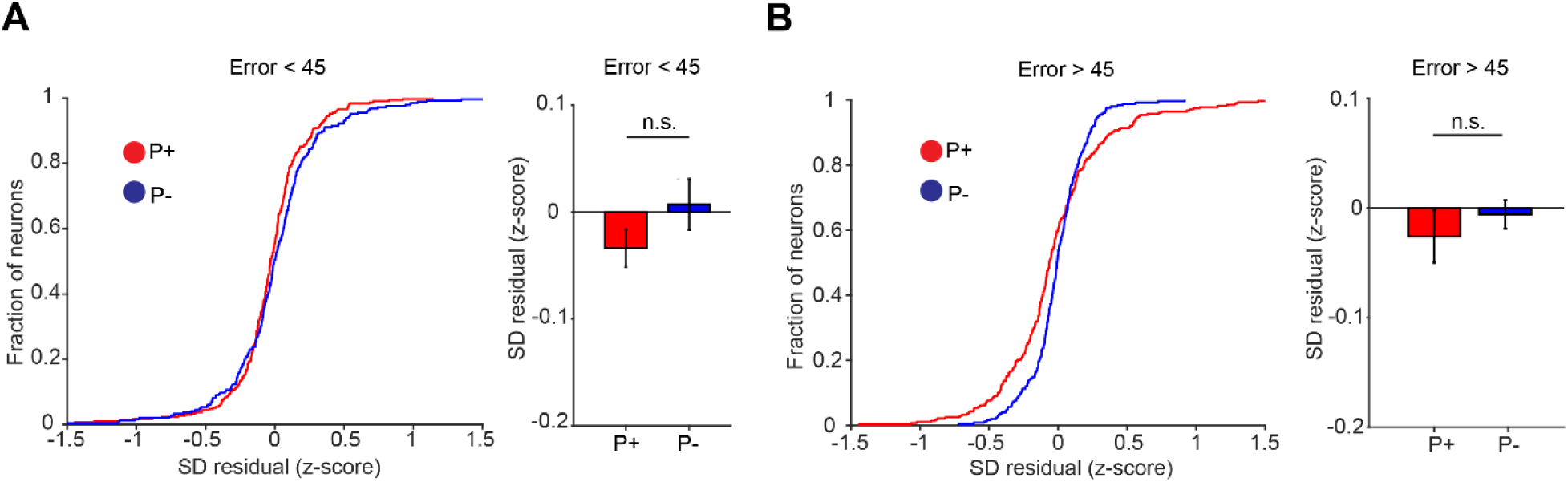
Dendritic signals do not represent absolute error values. a, The cumulative distribution function for SD residuals (z-scored per neuron) in P+ (red) and P-(blue) neurons during epochs where the absolute error was smaller than 45-degrees (t-test, p = 0.16; mean =-0.03 and 0.07; SEM = 0.017 and 0.02; n = 290 and 244 for P+ and P-neurons, respectively). **b**, Same as a, but for epochs in which absolute error was larger than 45-degrees (t-test, p = 0.47; mean-0.03 and-0.006; SEM = 0.02 and 0.01; n = 278 and 249 for P+ and P-neurons, respectively).

